# DOG1 prevents AFPs activation by AHG1 to control dormancy separately from ABA core signaling

**DOI:** 10.1101/2024.07.17.603870

**Authors:** Thorben Krüger, Dennis Brandt, Johanna Sodenkamp, Michael Gasper, Maida Romera-Branchat, Florian Ahloumessou, Elena Gehring, Julia Drotleff, Christopher Bell, Katharina Kramer, Jürgen Eirich, Wim J J Soppe, Iris Finkemeier, Guillaume Née

## Abstract

Seed dormancy determines germination timing and thereby critically influences seed plant adaptation and overall fitness. DELAY OF GERMINATION1 (DOG1) is a conserved central regulator of dormancy acting in concert with the phytohormone abscisic acid (ABA) through negative regulation of ABA HYPERSENSITIVE GERMINATION (AHG) 1 and AHG3 phosphatases. The current molecular mechanism of DOG1 signaling proposes that it regulates the activation state of central ABA-related SnRK2 kinases. Here, we unveil DOG1’s functional autonomy from the regulation of ABA core signaling components and unravel its pivotal control over the activation of ABSCISIC ACID INSENSITIVE FIVE BINDING PROTEINs (AFPs). Our data revealed a DOG1-AHG1-AFP relay in which AFPs’ genuine activation by AHG1 is contained by DOG1 to prevent the breakdown of maturation-imposed ABA responses independently of ABA-related kinase activation status. This work offers a molecular understanding of how plants fine-tune germination timing, while preserving seed responsiveness to adverse environmental cues, and thus represents a milestone in the realm of conservation and breeding programs.

**One-Sentence Summary:** Autonomous control of maturation-imposed ABA responses by DOG1 enables seeds to regulate dormancy and stress-reactivity traits independently.

## Main Text

With its remarkable resilience features, the seed represents a pivotal evolutionary innovation of plants fostering colonization of terrestrial areas (*1, 2*). The commitment to germination signifies the conclusion of the resilience phase thus, precise control of this process is paramount for success. Unfavorable environmental conditions during seed imbibition prevent germination (*3*). In addition, dormancy *sensu stricto*, an evolutionary and adaptative trait, restrains the sprouting of viable seeds, even under optimal conditions (*4*). Primary dormancy is established during seed maturation and is gradually released during dry storage (so-called after-ripening) or by environmental *stimuli* at imbibition (*5*). It ensures seedling establishment in the proper season, soil seed bank maintenance, promotes dispersal, and contributes to germination betting strategies (*6*).

The plant hormone abscisic acid (ABA), acting as a general growth inhibitor, is involved in many plant developmental processes and stress responses (*7*). Plants defective in ABA synthesis, perception, or core signaling transduction processes produce non-dormant seeds that cannot restrict their germination under stressful conditions (*8–11*). Degradation of ABA during imbibition promotes sprouting, while its *de novo* synthesis is required to prevent germination under stress or in dormant seeds (*12–16*). Hence, ABA is essential for primary seed dormancy induction during maturation as well as during imbibition to inhibit germination of dormant or non-dormant seeds under optimal or stress conditions, respectively. Dormancy loss slightly enlarges the environmental window permitting germination, but seeds’ stress-reactiveness is largely preserved indicating this trait to be distinct from stress responses (*17*). Dormancy alleviation mechanisms occurring in dry seeds are elusive but recognized to interact with the ABA-dependent germination arrest program upon imbibition (*18*). While both traits are undoubtedly dependent on ABA, the molecular framework by which seeds accommodate dormancy release while maintaining their capacity to stop germination if cues are met is not yet understood. Genetic determinants involved in the dormancy alleviation mechanisms during dry storage were identified by studying Arabidopsis natural variation in the length of after-ripening time required for germination (*19*). The gene underlying the major Quantitative Trait *Loci* effect, *DELAY OF GERMINATION 1* (*DOG1*), was shown to be required for seed dormancy and central for the local adaption of this trait to environmental conditions (*20–25*). Although DOG1 and ABA are both required for dormancy, these master regulators do not operate in a linear genetic pathway (*13, 25*).

ABA core signaling pathway consists of REGULATORY COMPONENT OF ABA RECEPTORs (RCARs), which bind ABA and consequently inhibit clade A PP2C phosphatases (PP2CAs) to prevent inhibitory dephosphorylation of class III SNF1-RELATED PROTEIN KINASEs of group 2 (SnRK2IIIs). Phosphorylation of SnRK2IIIs substrates downstream of the ABA core signaling leads to the activation of physiological ABA responses (*7*). *DOG1* encodes a protein of unknown molecular function (*25*). We previously found that DOG1 controls dormancy by physically interacting with ABA HYPERSENSITIVE GERMINATION 1 (AHG1) and AHG3 to negatively regulate their functions *in vivo* (*26*). AHG1, an atypical PP2CA, is an exceptional feature of the seed ABA signalosome and refractory to RCAR inhibition but negatively regulates ABA responses *in vivo,* like most PP2CAs (*27, 28*). Its genuine control by DOG1 explains the requirement of both DOG1 and ABA for dormancy (*26*). An independent report confirmed the existence of the DOG1-PP2C module and further proposed a direct inhibition of AHG1’s catalytic activity by DOG1 to maintain SnRK2IIIs phosphorylated, thereby active (*29*). SnRK2IIIs regulation by DOG1 would imply dormancy alleviation to be linked with desensitization to adverse conditions, thus disabling competitive advantages conferred by the seed habit. Noteworthy, null mutations of *DOG1* abolish dormancy but have minor effects on the seed sensitivity to ABA (*25, 26*). Hence, the DOG1-ABA molecular intertwining in the control of dormancy remains unresolved.

Using a proteomic-genetic integrated approach, we demonstrate that the DOG1-PP2C module does not participate in the control of SnRK2IIIs activity *in vivo* and, thus, that DOG1 controls dormancy independently of the ABA core signalosome activation. We identified a conserved phosphorylation site in two ABSCISIC ACID INSENSITIVE FIVE BINDING PROTEINs (AFPs) decreased in abundance in *dog1-1* mutant seeds and show that these are substrates of AHG1. We further show that AFPs are essential for the DOG1-PP2C module to control dormancy. Finally, through the combination of mutant alleles at different steps of the proposed pathway, we bring a compelling case that validates *in planta* a DOG1 signaling pathway controlling ABA responses independently of SnRK2IIIs activation status to oversee the dormancy trait. Our findings elucidate at the molecular level, how seeds adjust the timing of germination with minimal impact on their capacity to prevent sprouting under unfavorable conditions.

### DOG1 shapes the dry and early imbibed seed proteome and phosphoproteome

To identify downstream targets controlled by the DOG1-PP2C module *in vivo*, we performed a quantitative mass spectrometry analysis comparing the proteome and phosphoproteome of freshly harvested dormant NIL-DOG1 with non-dormant *dog1-1* seeds in dry and 6 hours imbibed seeds (Fig. 1A,B, and Supplementary Table 1). The *dog1-1* mutation affects 10.8 % (dry) and 13.1 % (imbibed) of the quantified protein groups and induced alterations in the seed phosphoproteome, with 19.9 % (dry) and 8.6 % (imbibed) of the quantified phospho-peptides (p-peptides) being significantly changed in abundance in mutant compared to NIL-DOG1 seeds (Fig. 1C-F). Coherently with the predisposition of *dog1-1* seeds to germinate at harvest, upregulated proteins were enriched in active metabolism, while downregulated proteins were enriched in stress and seed dormancy-associated GO terms (Fig. 1G). Proteins with regulated p-peptides were enriched in the GO terms cytosol and nucleus, as well as in RNA splicing, and ABA responses (Fig. 1H). This aligns with DOG1’s role in regulating the function of AHG1 and AHG3.

**Figure 1:**
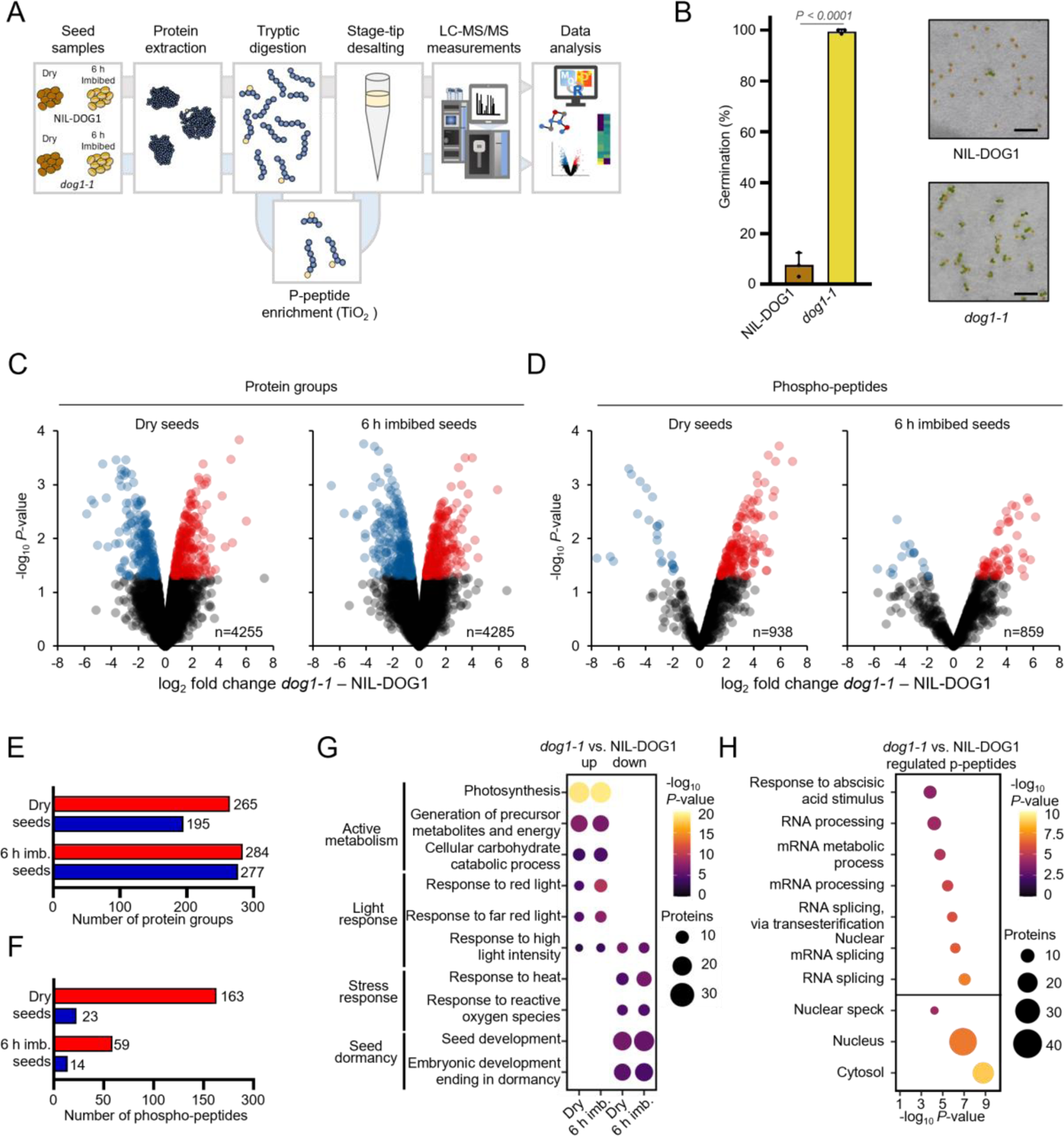
Proteome and phospho-proteome analyses of freshly harvested NIL-DOG1 and *dog1-1* seeds in the dry and 6 h water imbibed state. (**A**) Schematic workflow of the proteome analysis. Peptides for proteome and phospho-proteome analyses were prepared from the same seed extract and analyzed in parallel. (**B**) Germination capacity of NIL-DOG1 and *dog1-1* seeds at the sampling time of the proteomic experiments (Means±SD, n=3 biological replicates, unpaired t-test *P*-value is indicated). Scale bars in the picture represent 2 mm. (**C**&**D**) Volcano plots showing the regulation of (**C**) protein group abundance or (**D**) phospho-peptide abundance in dry (left panel) and 6 h imbibed (right panel) seeds of *dog1-1* compared to NIL-DOG1. (**E**) Number of protein groups and (**F**) phospho-peptides with significant changes (*P*-value<0.05) in abundance between genotypes. In **C-F**, down- and upregulation are shown in blue and red, respectively. (**G**) Gene Ontology (GO) enrichment for biological processes in dry or imbibed (imb.) seeds of proteins significantly regulated between genotypes. (**H**) GO biological process (top) and cellular component enrichment (bottom) of all (up- & down-regulated) phospho-proteins with significantly regulated phospho-peptides abundances either in dry, imbibed, or in both conditions.

### DOG1 stabilizes ABA responses during maturation and imbibition

To investigate the status of ABA responses in *dog1-1* seeds, we mined our proteomic data and found in either or both dry and imbibed seeds significantly regulated proteins annotated with the GO term “response to ABA”. This included well-known markers for positive ABA responses such as EM1, EM6, and RD29B that we found downregulated in *dog1-1* compared to NIL-DOG1 (fig. S1A,B). We further compared significant regulation between genotypes for all ABA-responsive proteins with changes observed at the mRNA levels in mutants defective in ABA synthesis and catabolism (*30*). We found that the regulations of ABA-responsive proteins in *dog1-1* are opposite to the effect of endogenous ABA (fig. S1C,D). When further analyzing ABI5 protein levels, we observed a significantly impaired accumulation in *dog1-1* compared to NIL-DOG1 dry and imbibed seeds (fig. S1E,F). Hence, our data show that during the end of maturation and early imbibition of freshly harvested seeds, DOG1 is required to properly establish ABA responses at the proteome level.

### DOG1 does not prevent SnRK2IIIs inhibition *in vivo*

P-peptides from 14 proteins implied in ABA responses, including 9 (cyto-)nuclear phosphoproteins, were significantly changed in abundance between genotypes in dry or imbibed seeds (Supplementary Table 2). As a hallmark, we found a p-peptide from SnRK2IIIs activation loop, modified at a position (i.e. S171 in SnRK2.6) directly associated with its ABA-dependent-upstream activation (*31–34*), to be more abundant in *dog1-1* compared to NIL-DOG dry seeds (Fig. 2A-C). We examined SnRK2IIIs accumulation in dry seeds and found protein levels to be similar in both genotypes (Fig. 2D). The normalization of p-peptide regulation on protein abundance indicated the activation mark residue of SnRK2IIIs to be approximately four times more phosphorylated in *dog1-1* compared to NIL-DOG1 (Fig. 2E). To further address the effect of DOG1 on ABA core signaling kinase activation and activity, we generated transgenic lines expressing a YFP:SnRK2.6 fusion protein in both NIL-DOG1 and *dog1-1* backgrounds (fig. S2A,B). The fusion construct localized in the nucleus and cytosol of 6 h and 24 h imbibed seeds embryos independently of the genetic background (fig. S2C). We used targeted proteomics to examine the modification status by phosphorylation of both S171 and S175/T176 after immunoprecipitation of the fusion protein from dry seed extracts of both backgrounds (fig. S3A,B). We found these residues to be more phosphorylated in *dog1-1* compared to NIL-DOG1 (fig. S3C-E). Next, we compared the kinase activity of YFP:SnRK2.6 purified from dry seeds of the two backgrounds and found that the kinase was more than two times more active when extracted from *dog1-1* compared to NIL-DOG1 seeds, aligning with their respective activation status (Fig. 2F). Altogether, our data directly demonstrate that *in vivo* DOG1’s function is not to prevent the deactivation of SnRK2IIIs as loss of *DOG1* function rather promotes their activity. Two p-peptides from group A bZIP C1 and C2 domains, were found to be more abundant in *dog1-1* compared to NIL-DOG1 seeds (fig. S4). This aligns with a higher SnRK2III activity in mutant seeds (*35*). Efficient SnRK2IIIs inhibition involved dephosphorylation of their activation loops and insertion into their catalytic clefts of a tryptophan conserved in PP2CAs and so-called “Trp-lock” (*36*). This residue is not conserved in AHG1, and its monocotyledon orthologues (fig. S5A,B). Molecular modeling suggests AHG1 to be unable to occupy SnRKIIIs active site compared to canonical PP2CAs (fig. S5C). To address if the enhanced SnRK2IIIs activation in *dog1-1* was associated with an increased sensitization to ABA, we analyzed transcript levels of the 14 Arabidopsis ABA receptors in dry NIL-DOG1 and *dog1-1* seeds (Fig. 2G). We found that the expression of seven *RCARs* was significantly upregulated in *dog1-1* (up to 7 times), while three receptors were downregulated (maximum two-fold). Altogether, our results indicate that the ABA core signaling pathway activity is primed in *dog1-1* seeds. Yet, this activation does not override the non-dormant phenotype of *dog1-1* seeds. Consequently, the DOG1-PP2C module must operate *via* a molecular mechanism distinct from the regulation of SnRK2IIIs activation.

**Figure 2:**
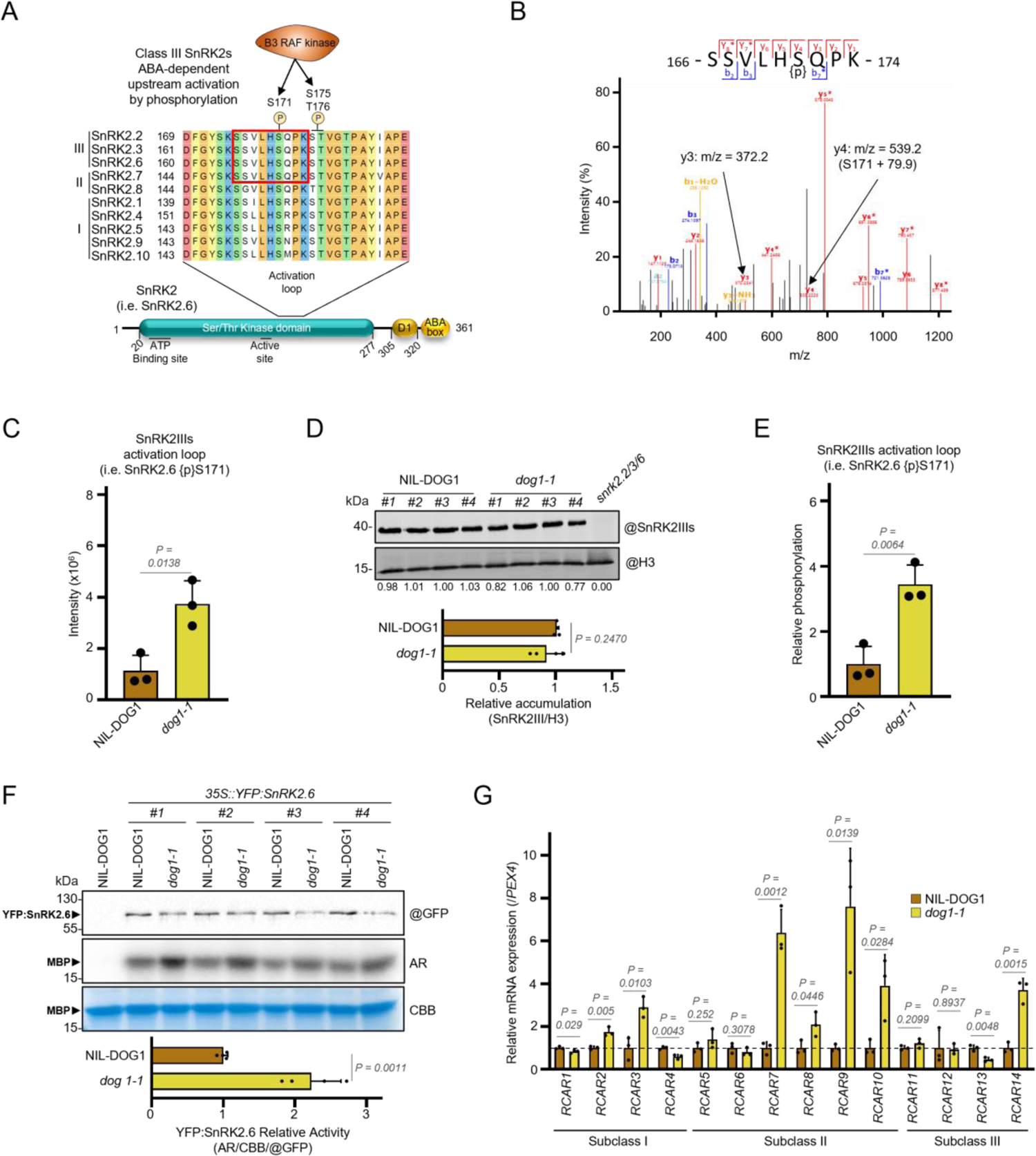
ABA core signaling is primed in *dog1-1* seeds. **(A)** Schematic representation of SnRK2IIIs protein (using SnRK2.6 as an example), and of amino acid sequence alignment of Arabidopsis SnRK2s’ activation loops. The tryptic p-peptide more abundant in *dog1-1* compared to NIL-DOG1 is highlighted in red. Phosphorylation of SnRK2III activation loop is mediated by RAF kinases. (**B**) MS^2^ spectrum of the tryptic peptide phosphorylated at S171 (in SnRK2.6). (**C**) MS^1^ intensities of the SnRK2IIIs activation loop p-peptide in dry NIL-DOG1 and *dog1-1* seeds (Means±SD, n=3 biological replicates). (**D**) Western blot quantification of SnRK2IIIs in dry NIL-DOG1 and *dog1-1* seeds. The bar chart shows protein SnRK2III accumulation normalized to the Histone 3 (H3) loading control (Means±SD, n=4 biological replicates). (**E**) Phosphorylation of SnRK2IIIs activation loop between dry NIL-DOG1 and *dog1-1* seeds after normalization of peptide intensity to the protein abundance change (Means±SD, n=3 biological replicates). (**F**) Kinase activity of YFP-SnRK2.6 immunoprecipitated from dry seed of NIL-DOG1 and *dog1-1* background. YFP-SnR2.6 bound beads were used to phosphorylate MBP *in vitro* in the presence of ^32^P γ-ATP. The phosphorylation of MBP with radiolabeled phosphate was assessed using autoradiography (AR), loading of MBP evaluated by Coomassie Brilliant Blue (CBB) staining and the amount of YFP:SnRK2.6 investigated with anti-GFP. Signal intensities of phosphorylated MBP (autoradiography/AR), total MBP (Coomassie Brilliant Blue/CBB) and YFP-SnRK2.6 (@GFP) amounts (bands marked with arrows) were determined by densitometric analyses. The bar chart shows the normalized (AR/CBB/@GFP) kinase activity (Means±SD, n=4 biological replicates). IP from non-transgenic NIL-DOG1 protein extract served as negative control. (**G**) Expression of the 14 Arabidopsis ABA receptors from the *RCAR* family in dry seeds of NIL-DOG1(Means±SD, n=3 biological replicates). The data presented in **C-G** show levels relative to NIL-DOG1 (arbitrary mean value of 1) and unpaired t-test *P*-values are indicated. In **D** and **F**, the # indicate independent biological replicates.

### The redundant negative regulators of seed dormancy, AFP1 and AFP2, are less phosphorylated in *dog1-1*

Since DOG1 negatively regulates AHG1 and AHG3 phosphatases in the nucleus, we searched for nuclear proteins that were less phosphorylated in *dog1-1* mutants. Our analysis identified p-peptides deriving from AFP1 and AFP2 modified at S115 and S112, respectively (fig. S6A,B). This residue is conserved in all four Arabidopsis AFPs and located in their B-domain (fig. S6C). The AFP1 p-peptide was found to be eight times less abundant in dry seeds of *dog1-1* compared to NIL-DOG1 (Fig. 3A). Protein quantification revealed AFP1 abundance to be reduced by 50 % in *dog1-1* dry seed (Fig. 3B). The normalization of the p-peptide change relative to the protein change revealed that the AFP1 protein pool was about four-times less phosphorylated at S115 compared to NIL-DOG1 (Fig. 3C). Our data also indicated that the AFP2 p-peptide was less abundant in imbibed mutant seeds compared to NIL-DOG1 (Supplementary Table 2). Similar to *DOG1* and *AHG1*, the expression of *AFPs* is mainly restricted to seed tissues (fig. S7A). As the maternal environment strongly influences primary seed dormancy levels in Arabidopsis (*18*), we examined if *AFP* expression is responsive to temperature *stimuli* during seed maturation, resulting in contrasting dormancy at harvest (fig. S7B,C). The expression of all four *AFPs* was significantly lower in dormant compared to non-dormant seed batches (fig. S7D), indicating an association between dormancy levels and *AFP* expression. To address the importance of the different AFPs in the control of seed dormancy *sensu stricto*, we characterized this trait for single and multiple *afp* mutants. Our analysis demonstrated that AFP1, AFP2, and AFP3 act as partially redundant negative regulators of dormancy (fig. S8A). The hyper-dormant phenotype of the triple *afp1-5/2-2/3-1* mutant was still observable in seed batches obtained from conditions resulting in shallow dormancy of Col-0 seeds at harvest (fig. S8B). We did not observe an enhancement of the *afp2-2* phenotype by the *afp4-2* mutant allele (fig. S8C). Potent ectopic overexpression of *AFPs* leads to severe maturation defects and compromises the viability of homozygous *35S::YFP:AFP2* seeds (*37, 38*). Thus, we analyzed the dormancy of seed progenies from *35S::YFP:AFP2* (+/-) hemizygous parental lines. Despite matured in a dormancy-promoting maternal environment, we found that transgenic seeds were fully non-dormant compared to Col-0 (fig. S9A,B), confirming that AFPs are negative regulators of seed dormancy. To further address the importance of ABA level dynamics for AFPs to control germination, we used fluridone or abscinazole-E3M (ABZ-E3M) to prevent ABA *de novo* synthesis or catabolism, respectively. While fluridone effectively promoted the germination of freshly harvested Col-0 seeds, the *afp1-5/2-2/3-1* triple mutant seeds were not responsive to this treatment (fig. S8D). Abscinazole-E3M significantly reduced germination of after ripened Col-0 but not *35S::YFP:AFP2* (+/-) seeds progenies (fig. S9C). Thus, the function of AFP in the control of germination appears independent of ABA metabolism during imbibition. These findings prompted us to investigate the involvement of AFPs in the regulation of seed dormancy by the DOG1-PP2C module.

**Figure 3:**
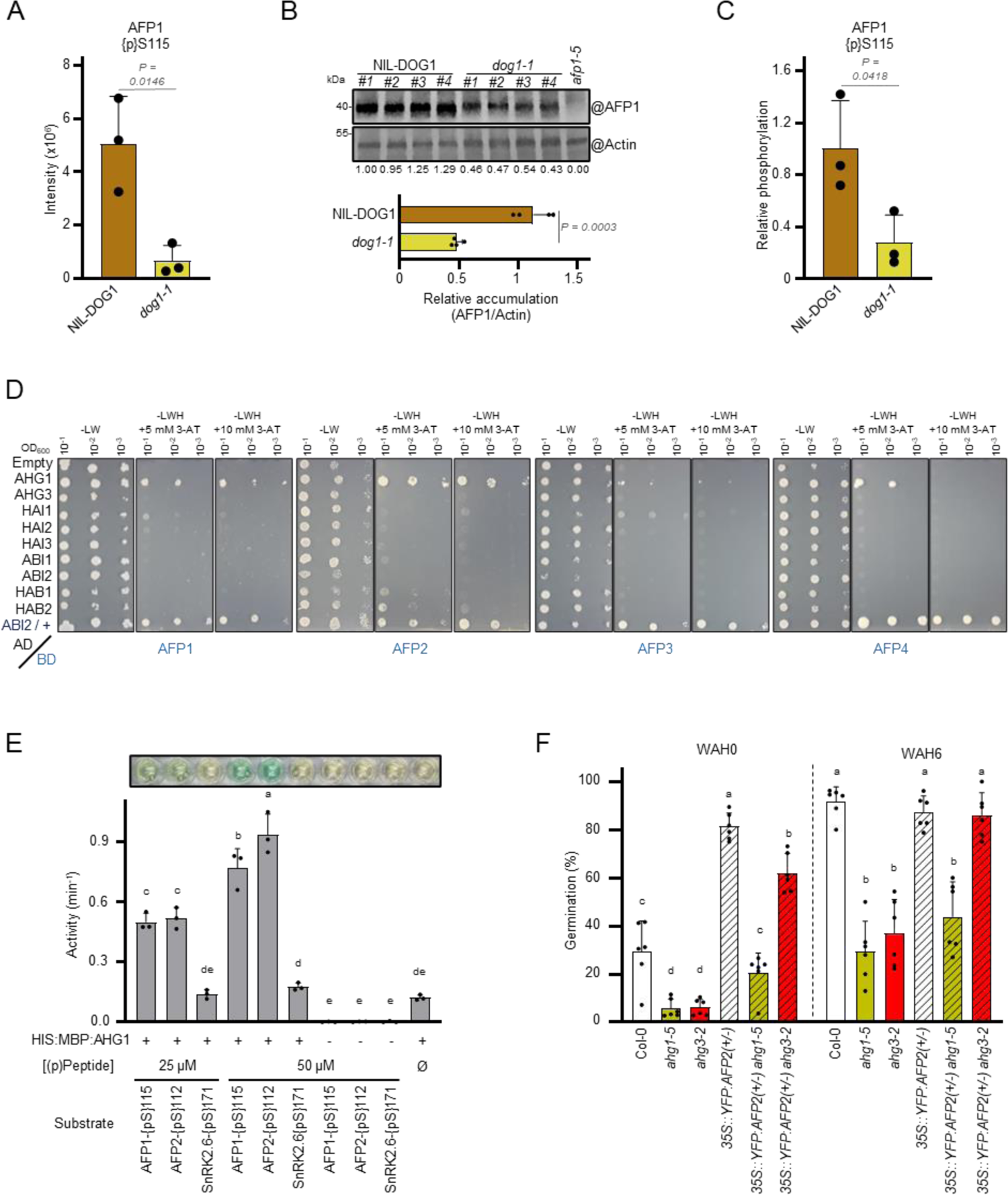
AFPs are substrates of AHG1, and AFP2 functionally requires AHG1 to control germination *in vivo*. (**A**) MS intensity of AFP1 p-peptides containing {pS}115 (Means±SD, n=3 biological replicates). (**B**) Western blot quantification of AFP1 in dry NIL-DOG1 and *dog1-1* seeds. The bar chart shows protein accumulation of AFP1 normalized to the Actin loading control (Means±SD, n=4 biological replicates). (**C**) Phosphorylation of AFP1 {pS}115 between dry NIL-DOG1 and *dog1-1* seeds after normalization of peptide intensity to protein abundance change (Means±SD, n=3 biological replicates). (**D**) Family-wide yeast-2-hybrid screen between Arabidopsis clade A PP2Cs and AFPs. Control medium lacking leucine and tryptophan =-LW; interaction medium additionally lacking histidine = -LWH; 3-AT = 3-aminotriazole. Interaction of ABI2 with SnRK2.6 (+) served as positive control. Negative control assays for BD-AFPs self-activation are shown in fig. S10. (**E**) *In vitro* phosphatase assays using HIS:MBP:AHG1 and synthetic phosphorylated peptides of AFP1{pS}115, AFP2{pS}112 and SnRK2IIIs activation loop (i.e. SnRK2.6{pS}171). Assays in which the substrate or the enzyme was omitted served as negative controls (Means±SD, n=3 independent assays). The top panel shows a representative picture of the colorimetric assay result. A positive control for SnRK2.6{pS}171 dephosphorylation is presented in fig. S12. (**F**) Germination capacity of seed progenies from *35S::YFP:AFP2*(+/-) in Col-0, *ahg1-5* or *ahg3-2* mutant backgrounds compared to their parental lines at harvest and after 6 weeks after harvest (WAH) (Means±SD, n=6 biological replicates). The data presented in **A-C** show levels relative to NIL-DOG1 (arbitrary mean value of 1) and unpaired t-test *P*-values are indicated. In **B**, the # indicates independent biological replicates. In **E** and **F**, letters on the top of the bars indicate significantly different groups using a one-way ANOVA with a Tukey HSD test (α=0.05).

### AFPs are genuine substrates of AHG1

To determine if AFPs are substrates of PP2CAs, we investigated the physical protein interactions between the four AFPs and the nine PP2CAs of Arabidopsis in a yeast two-hybrid system. We observed that all AFPs exclusively interacted with AHG1 (Fig. 3D and fig. S10). We confirmed the specificity of the AFP-AHG1 interactions *in planta* by conducting co-immunoprecipitation (Co-IP) assays using *Nicotiana benthamiana* leaves transiently co-expressing *Venus:AFPs* and *3xHA:PP2CAs* (fig. S11). Lastly, we used a recombinant protein with assessed quality and activity to examine AHG1’s *in vitro* capability to directly dephosphorylate AFP1 and AFP2 phosphoserines {pS} (fig. S12A-C). We found that AFP1{pS}115 and AFP2{pS}112 are substrates of AHG1 (Fig. 3E). In contrast, AHG1 displayed no activity towards the SnRK2IIIs phosphoserine, which was hyperphosphorylated in *dog1-1* (Fig. 3E and fig. S12D).

### AFP2 requires AHG1 to control germination

To investigate the requirement of AHG1 for AFP2 functions *in vivo*, we crossed the *AFP2* over-expression line with *ahg* mutants and isolated hemizygous *35S::YFP:AFP2* (+/-) plants in homozygous *ahg1-5* and *ahg3-2* backgrounds. While *YFP:AFP2* overexpression abolished dormancy in Col-0, we observed that seed progenies from *35S::YFP:AFP2* (+/-) in *ahg1-5* germinated to a lower extent compared to Col-0 seeds (Fig. 3F). These data demonstrate that AHG1 is genetically required for AFP2 to function in the control of dormancy. The seed progenies from *35S::YFP:AFP2* (+/-) in the *ahg3-2* background exhibited only a marginal increase in dormancy compared to the seed batches from *35S::YFP:AFP2* (+/-) in the Col-0 background. This indicates that AFP2’s function during germination is not dependent on AHG3, which is in line with the specific interactions of AFPs with AHG1.

### The non-dormant phenotype of *dog1* seeds is partially dependent on AFP1 and AFP2

To investigate the genetic relationship between DOG1 and AFPs, we generated combinatorial *afp* and *dog1-2* mutants. Germination analysis of double to quadruple mutant seeds showed that in contrast to *dog1-2*, the triple *afp1-5/2-2 dog1-2* mutant produces about 25 % of dormant seeds. The additional mutation of *afp3-1* in the quadruple mutant did not further enhance the dormancy levels in the context of the *dog1-2* background (Fig. 4A). No effect of the *afp4-2* allele was observed when comparing seed dormancy of double *afp2-2 dog1-2* and triple *afp2-2/4-2 dog1-2* mutants (fig. S13A). We reproduced our results using independent *afp* mutant alleles (fig. S13B). Hence, *afp1* and *afp2* are sufficient to reintroduce dormancy in *dog1-2,* indicating that DOG1 signaling requires both AFP1 and AFP2.

**Figure 4:**
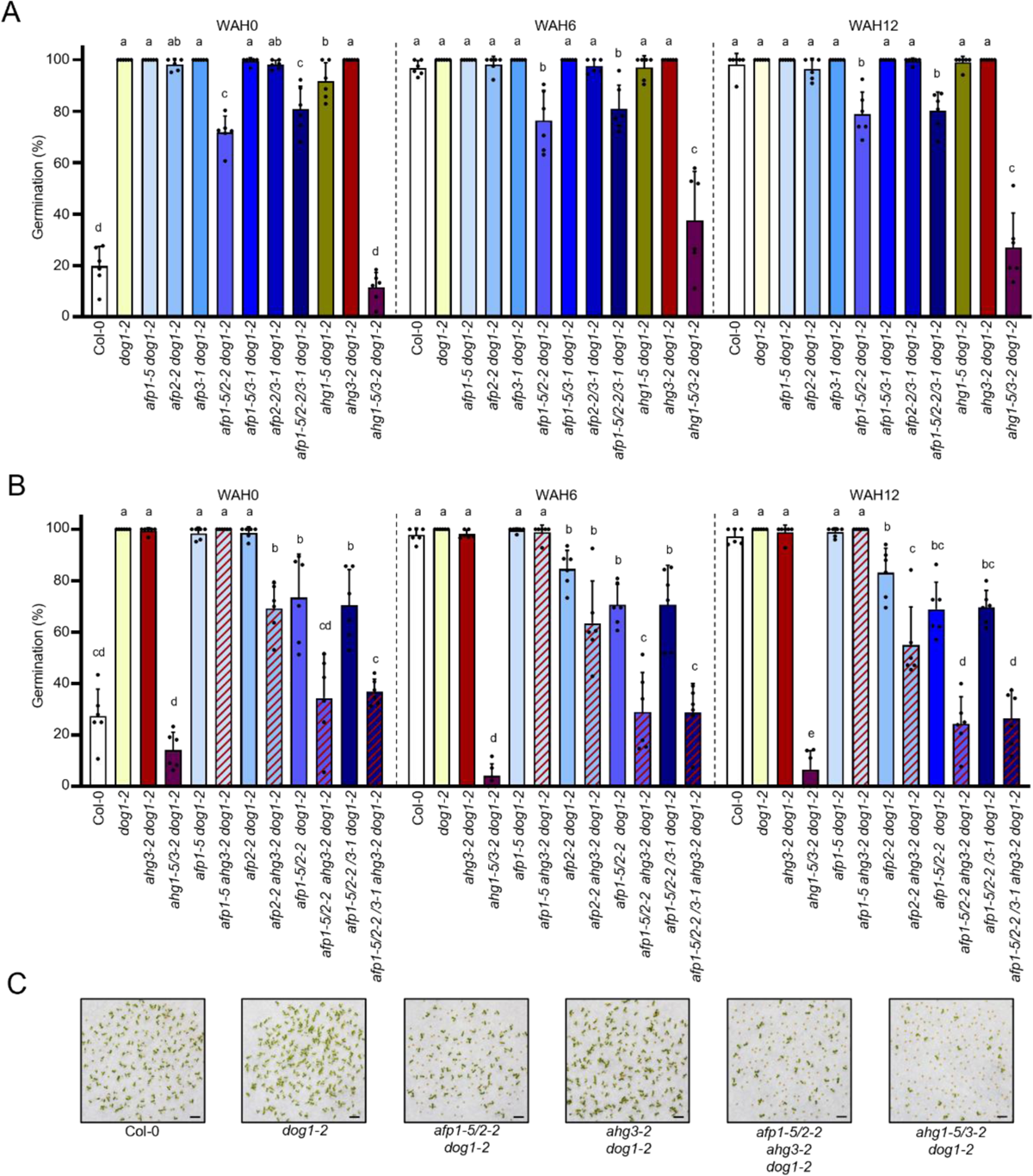
AFPs are required by the DOG1-PP2C module and operate specifically downstream of AHG1 in the control of seed dormancy. (**A**) Germination capacity of single and multiple *afps* mutants in a *dog1-2* or, (**B**) an *ahg3-2 dog1-2* genetic background at harvest, 6 and 12 weeks after harvest (WAH) (Means±SD, n=6 biological replicates). Letters on the top of the bars indicate significantly different groups using a one-way ANOVA with a Tukey HSD test (α=0.05). (**C**) Representative pictures showing the germination capacity of key combinatorial mutants after 32 WAH resuming the importance of AFP1 and AFP2 for the control of germination by the AHG1-related branch of the DOG1-PP2C module.

### AFPs function downstream of the AHG1 branch

Our previous work showed that DOG1 relies on both AHG1 and AHG3 functions *in planta* (*26*). Here, we have shown that AFPs interact specifically with AHG1 and that this atypical PP2CA is essential for the function of AFP2 in seed dormancy (Fig. 3D,F and fig. S11). Hence, the activation of AHG3 might be responsible for the approximately 75 % of germination in the *afp1-5 2-2 dog1-2* triple mutant seed batches. We tested this hypothesis by combining *ahg3-2* and *afp* mutants. We found seed dormancy of *afp2-2 ahg3-2*, *afp1-5/2-2 ahg3-2,* and *afp2-2/3-1 ahg3-2* to be highly enhanced compared to their relative parents with an *AHG3* Wild Type (WT) allele (fig. S14). Next, we analyzed the ability of *ahg3-2* to enhance seed dormancy of multiple *afp dog1-2* seeds. We observed that dormancy of the quadruple *afp1-5/2-2 ahg3-2 dog1-2* mutant was much stronger than the triple *afp1-5 2-2 dog1-2* or double *ahg3-2 dog1-2* mutant seeds at harvest (Fig. 4B,C). This shows that the AHG1 branch of the DOG1-PP2C module requires AFP1 and AFP2 to control seed dormancy. We noted that like the *ahg1-5/3-2 dog1-2* mutant, the dormancy level of quadruple *afp1-5/2-2 ahg3-2 dog1-2* or quintuple *afp1-5/2-2/3-1 ahg3-2 dog1-2* mutant seeds stayed stable even after extended storage (Fig. 4B,C). The insensitivity of these mutant genotypes to after-ripening indicates that they are unable to operate the molecular switch that normally permits growth after dormancy alleviation.

## Discussion

The role of ABA in inducing growth arrest during stress responses can be traced back to the earliest land plants, although prototypical components of the ABA core signaling pathway are already present in algae (*2*). ABA responses evolved to reach their maximum intricacy in seed plants where for example, ABA core signaling is encoded by multigenic families and embedded in a complex regulatory network. Such evolutive sophistication enabled the ubiquitous ABA signal to precisely control diverse physiological responses based on multiple developmental and environmental inputs (*28*). Dormancy is one of these phenomena. It is built up during maturation, but the resulting inhibition of sprouting takes place upon imbibition, accounting for differences in the germination level of seeds that have after-ripened or not. Although requiring ABA during seed maturation, dormancy depth is not directly correlated with ABA levels in dry seeds but is positively associated with DOG1 accumulation (*12, 13*). ABA and DOG1 signaling pathways crosstalk at the level of the PP2CAs in the control of seed dormancy (*26, 29*).

Here, we addressed the question whether or not DOG1 acts through modification of the canonical ABA core pathway, which is a long-standing debate in the seed biology and ABA signaling fields (*18, 28, 39, 40*). In agreement with previous transcriptome studies (*41*), we found that DOG1 is required for the proper establishment of ABA responses at the proteome level during seed maturation and early imbibition (Fig. 1 and fig S1). However, our data demonstrate that DOG1 does not play a role in preserving SnRK2IIIs in an active state *in vivo*. The phosphorylation of conserved SnRK2IIIs residues associated with their activation is enhanced in non-dormant *dog1-1* when compared to dormant NIL-DOG1 dry seeds (Fig. 2A-E and fig. S3). In agreement with enhanced activation marks, extractible SnRK2.6 activity from dry seed material was higher in *dog1-1* compared to NIL-DOG1 (Fig. 2F). The increased activation of the central ABA kinases was further supported by elevated levels of the two p-peptides corresponding to SnRK2IIIs-activated group A bZIPs in mutant dry *dog1-1* seeds (Fig. S4), despite the fact that these transcription factors are downregulated in this mutant (*41*) (fig. S1E). SnRK2IIIs deactivation follows a two-step mechanism whereby the conserved Trp-lock residue of PP2CAs is inserted into the kinase catalytic cleft after activation loop dephosphorylation to blunt basal activity and prevent auto- or trans-reactivation of SnRK2 isoforms (*36*). Absence of the Trp-lock residue is a key feature of AHG1 and its orthologs in monocots (fig. S5A,B). Molecular modeling suggests that the valine residue present at this position in AHG1 cannot occupy the kinase active site as the tryptophan of other PP2CAs (fig. S5C). This limitation might explain previous observations that AHG1 is less effective than other PP2CAs in suppressing SnRK2.6 activity *in vitro* and that AHG1 is incapable of impairing SnRK2-mediated ABA responses using an *in vivo* reconstituted system (*42–44*). ABA levels were previously shown to be lower in dry seeds of *dog1-1* compared to NIL-DOG1 (*13*), but consistently with an increased SnRKIIIs activation, we found an upregulation of ABA-receptors from subclass I and II (Fig. 2G), which inhibit canonical PP2CAs in the absence or at low concentrations of ABA (*44, 45*). Altogether, our data show that the ABA core signaling machinery is fully functional in *dog1-1* mutant seeds through, increase of sensitization, priming of SnRK2III activity, and activation of direct downstream effectors. Yet, this molecular priming does not result in stable ABA responses (fig. S1), nor does it override the non-dormant phenotype of *dog1-1* (Fig. 1B). Consequently, in stark contrast to the current model, our data demonstrates that DOG1 function is not to regulate SnRK2IIIs activation through the inhibition of AHG1 and thus that DOG1 controls dormancy independently from the central ABA kinase hub.

We noted that p-peptides from AFP1 and AFP2, with a conserved modification site, showed reduced abundances in *dog1-1* compared to NIL-DOG1 seeds (Supplementary Table 2). At this position, AFP1 exhibited approximately four times less phosphorylation in *dog1-1* compared to NIL-DOG1 dry seeds (Fig. 3C), prompting us to explore AFPs as critical targets of the DOG1-PP2C module to control dormancy. Only AFP2 was proposed to participate in dormancy so far (*46, 47*). Here, we show that AFP1, AFP2, and AFP3 function as genetically highly redundant negative regulators of seed dormancy *sensu stricto* since the cumulation of their mutant alleles gradually enhanced this trait (fig. S8A-C and S13B). AHG1 is the only PP2CA interacting with AFPs (Fig. 3D and fig. S11), and we showed that it directly targets the p-sites regulated in *dog1-1*: AFP1{pS}115 and AFP2 {pS}112 (Fig. 3E and fig. S12). We demonstrated that AFP2 requires AHG1 to release dormancy (Fig. 3F). Coherently with AFPs-AHG1 genuine interaction, we only found a marginal effect of the *ahg3* mutant allele on the dormancy phenotype of transgenic *AFP2* seeds (Fig. 3F). This suggests that AHG1 governs the upstream activation of AFPs in part through dephosphorylation of a conserved B-domain residue. An inhibitory role of phosphorylation on AFP is further supported by the fact that the kinase SnRK1α1 antagonizes the effect of transient *AFP2* expression in *Nicotiana benthamiana* leaves (*48*).

AFP activities are relevant for the DOG1 pathway since *afp1* and *afp2* mutations partially revert the *dog1* dormancy phenotype (Fig. 4A and fig. S13B). Analysis of higher-order combinatorial mutants including *afps*, *ahg3* and *dog1* loss-of-function alleles, demonstrated that the contribution of the *ahg1* allele in the highly dormant *ahg1-5/3-2 dog1-2* mutant can largely be substituted by *afp1* and *afp2* mutations (Fig. 4B,C). This result confirms AFP-independent functions for AHG3 and validates genetically that AFP1 and AFP2 are pivotal and genuine targets of AHG1 crucial for the DOG1-PP2C module to control seed dormancy. Pharmacological assays to manipulate ABA dynamics at imbibition suggest that AFP’s function in the control of germination is independent of ABA levels (fig. S8D and fig. S9C) This aligns with the reported extreme seed ABA insensitivity of constitutive *YFP:AFP2* overexpression lines used in this study (*37, 38*).

AFPs are proteins of unknown function. ABA promotes the accumulation of AFP1 and AFP2, but they rapidly decline through protein post-translational modification and proteasome-dependent mechanisms under stress-free conditions (*38, 46, 49*). Thus, they act as a rheostat transiently generated by ABA to moderate the magnitude of ABA responses or terminate them after stress relief (*50*). DOG1 protein remains stable in dry and imbibed seeds, regardless of the imbibition outcomes, but it is modified during dry storage and seed water uptake suggesting that DOG1 has lost its activity in after-ripened seeds (*13*). D*e novo* production of *DOG1* expression even as short as three hours after seed imbibition, thought transgenic inducible expression, does not prevent germination of freshly harvested *dog1-1* seeds (*13*). This supports the fact that the amount of active DOG1 protein readily present at early imbibition determines whether a seed is dormant or not. This indicates that the impact of DOG1 on the control of germination is limited to a narrow time window following water uptake and inefficient at later imbibitional stages of non-dormant seeds in agreement with its irrelevance in the process to stop germination if stress conditions are met during imbibition.

The DOG1-AHG1-AFPs molecular relay presented here significantly expands our understanding of the molecular tinkering of ABA responses that allow seeds to accommodate germination time and stress reactiveness. We propose a non-conflicting bi-modal molecular system for the mediation of ABA responses during seed imbibition by DOG1 and ABA (Fig. 5). DOG1 prevents AHG1 from promoting AFPs-mediated shut-down of maturation-imposed ABA responses, thereby maintaining dormancy regardless of the initial drop of ABA levels upon imbibition. Because the DOG1-PP2C module functions are separated from the regulation of SnRK2IIIs activity, it keeps ABA signaling primed independently of dormancy depth. This allows non-dormant seeds to re-induce ABA responses and arrest germination upon sensing stress quickly and cost-efficiently. This dual molecular system allows seeds to bypass trade-offs between growth promotion and stress tolerance and to precisely adjust their dormancy level with minimal impact on their ability to withstand adverse conditions.

**Figure 5:**
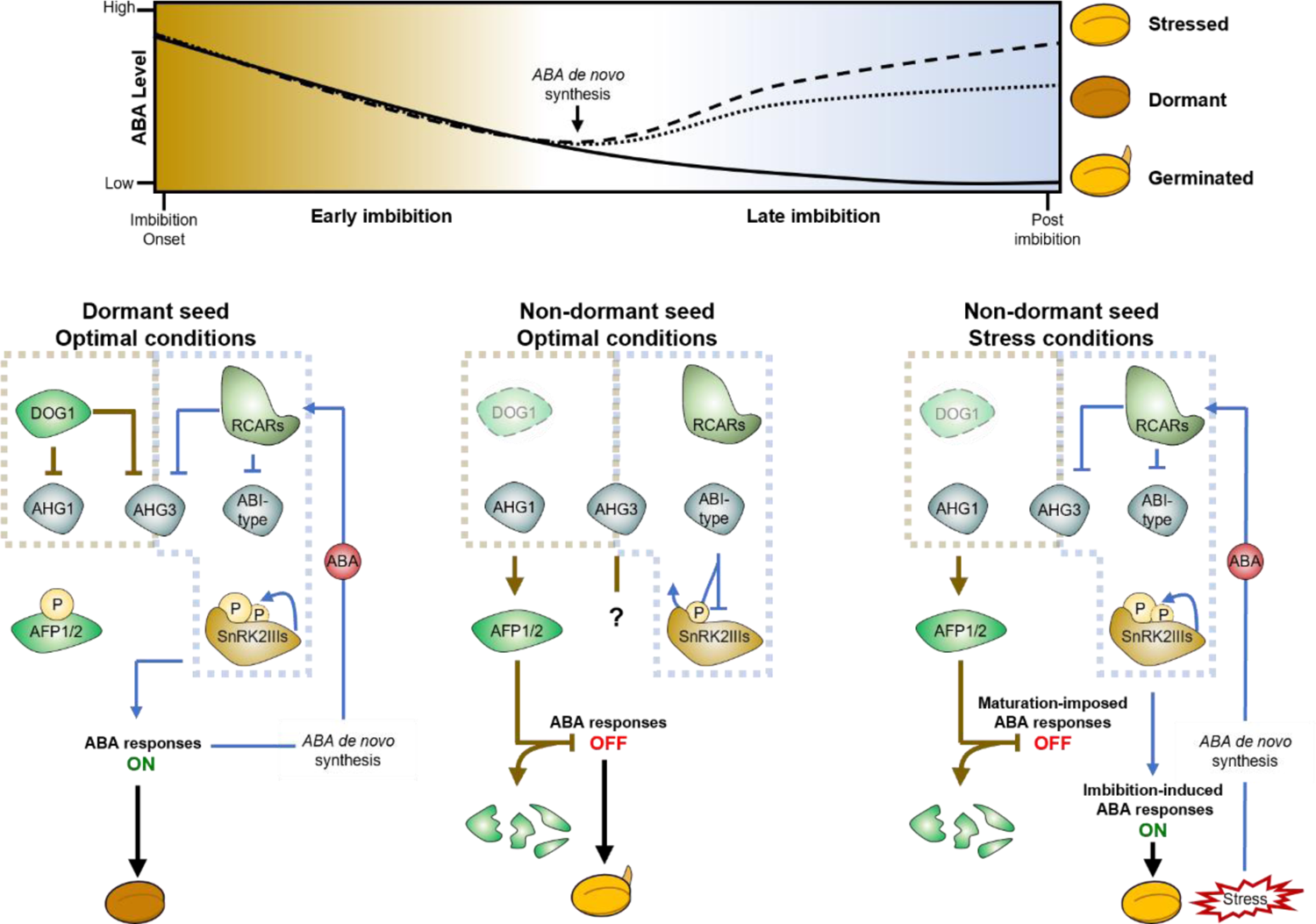
Proposed model for the accommodation of both dormancy and stress reactiveness during seed imbibition by a bi-modular control of ABA responses. Schematic representation of ABA level dynamics (top) and the operation of the DOG1-PP2C module and the ABA core signaling in dormant (bottom left), non-dormant unstressed (bottom center), or non-dormant stressed (bottom right) seeds during imbibition. ABA levels decrease upon imbibition regardless of the seed’s stress or dormancy status, but ABA is *de novo* synthesized at later imbibitional stages in non-germinated seeds (dormant or stressed) to prevent sprouting after long-term hydration. In dormant seeds imbibed under optimal conditions, DOG1 is active and prevents AFP activation by AHG1, thus maintaining ABA responses imposed during maturation, independently of the initial drop in ABA level during early imbibition. In the later stages, active seed maturation responses induce ABA *de novo* synthesis that, together with the priming of SnRK2IIIs activation sustain active ABA responses post imbibition and preclude germination. In non-dormant seed (e.g. after-ripened or *dog1* mutant seeds) imbibed under optimal conditions, where DOG1 is not active (dashed border; shaded), AHG1 and AHG3 functions are revived. AHG1 dephosphorylates AFP1 and AFP2, which switches off ABA responses imposed during maturation. The absence of stress during early imbibition leads to AFP disposal. Disrupted ABA responses and the absence of efficient ABA *de novo* synthesis under a stress-free environment promote germination and growth. In non-dormant seed imbibed under stress conditions, DOG1 is not active, leading to AFP activation and destabilization. While ABA responses imposed during maturation are terminated, SnRK2IIIs activation stays primed (in *dog1* mutants, the founder activating phosphorylation mark is preserved). High levels of ABA from stress-induced *de novo* synthesis at the later stages of imbibition prevent canonical PP2CA-mediated SnRK2III repression, allowing a rapid re-activation of the kinases to induce the ABA imbibitional responses triggering the arrest of germination. The question mark indicates the existence of unknown but AFP-independent AHG3 functions in the control of germination. Molecular events occurring during early or late seed imbibition appear as brown or blue connectors, respectively. The DOG1-PP2C module (brown) and the ABA core signaling pathway (blue) are boxed.

Germination timing is crucial to synchronize growth potential with seasonally-fluctuating ambient conditions, thus vital for the successful establishment of seed plants in nature. The initial stages of plant domestication witnessed a strategic shift in resource allocation, diverting focus from the “fitness package” towards enhancing yield. Notably, cereal domestication marked a pivotal moment with the deliberate selection of genotypes lacking dormancy, ensuring swift and uniform germination across expansive fields (*51*). Beyond the induction of major issues like preharvest sprouting and varying malting quality, the loss of genetic diversity associated with the domestication of dormancy has reduced global plant fitness through selective sweeps at and around *loci* involved in ABA metabolism, causing pleiotropic effects (*52–55*). This highlights the need to neo-domesticate the dormancy trait for the creation of “wild-relative-like” genetic restorer lines that maintain high fitness while keeping germination characteristics adapted to our food production system. DOG1 function is conserved in higher plants, the existence of the DOG1-PP2C module has been documented in crop species, and AFP function in the control of dormancy has been reported in wheat (TaAFP-B) and rice (Mediator of OsbZIP46 Deactivation and Degradation / OsMODD) (*56–61*). Importantly, the genetic edition of OsMODD has recently been shown promising for the generation of rice restorer lines (*61*). The typical sequence feature of AHG1 and the modification of AFP the B-domain conserved serine residues by phosphorylation we described in Arabidopsis also exist in these major crop species (fig. S5A,B and fig. S6D). While the evolutionary origin of the DOG1 signaling pathway presented herein is still to be defined, it appears largely conserved in seed plants. Through the mapping of a pathway controlling dormancy independently of other ABA-related germination traits, our findings offer so far unexplored opportunities for breeding programs aiming to deliver chemical-free solutions for contemporary environmental and food security challenges.

## Acknowledgments

We thank Prof. Ruth Finkelstein (University of California, Santa Barbara) for providing the *35S::YFP:AFP2* (+/-) transgenic and *abi5-7* mutant lines and Prof. Luis Lopez-Molina (University of Geneva) for providing the AFP1 antibody. We thank Dr. Guillaume Brun and Dr. Stefan Weinl (University of Muenster) for providing an aliquot of ABZ-E3M and support for isotope related work respectively. We also thank Anne Harzen (MPI Cologne) for MS sample preparation and Emelie Felgenhauer and Thomas Willenborg (University of Muenster) for their technical support during seed batch production/processing and mutant genotyping. We thank Paulina Heinkow (University of Münster) for technical assistance, and for maintaining the LC-MS/MS instruments at the MSPUB. We dedicate this work to our friend and former colleague Florian Ahloumessou who passed away on the 04^th^ August 2019 at the age of 28.

## Funding

This work was supported by the Deutsche Forschungsgemeinschaft Grant# NE2296 to GN and grant number 469950637 to IF as well as INST 211/744-1 FUGG for instrumentation. In addition, this work was supported by funding from the Max-Planck Society (IF, KK, GN and WS.). TK was supported by a fellowship of the Studienstiftung des deutschen Volkes and by the department of biology of the University of Muenster. The research stay of FA at the University of Muenster was funded by the DAAD. Open access funding was provided by the University of Muenster.

## Author contributions

G.N. conceived the study, applied for funding, and designed the experiments. T.K. and G.N. performed most of the experiments with the help of D.B. for Y2H assay, C.B. for Co-IP assays, J.S, E.G., F.A, M.G., J.D and, D.B during to mutant isolation and phenotyping. D.B., J.S., M.G., J.D., E.G., C.B. conducted their experiments as integral components of their degree programs at the University of Muenster. M.R.B. performed transcript analysis. I.F., J.E and K.K provided MS measurements and support for method development, Max Quant and Perseus analyses. I.F. provided funding support and supervision. G.N. supervised the project, analyzed the data with the help of T.K., and drafted the manuscript. G.N. wrote the final version, with edits from T.K., W.S., and I.F.

## Competing interests

Authors declare that they have no competing interests.

## Data and materials availability

Mass spectrometry proteomics data have been deposited in the ProteomeXchange Consortium (http://proteomecentral.proteomexchange.org) via the JPOSTDB partner repository with the data set identifier PXD046985 and PXD053457. Sequence data from this article can be found in The Arabidopsis Information Resource-TAIR database (www.arabidopsis.org) under the following accession numbers: AFP1 (AT1G69260), AFP2 (AT1G13740), AFP3 (AT3G29575), AFP4 (AT3G02140), AHG1 (AT5G51760), AHG3 (AT3G11410), ABI1 (AT4G26080), ABI2 (AT5G57050), HAB1 (AT1G72770), HAB2 (AT1G17550), HAI1 (AT5G59220), HAI2 (AT1G07430), HAI3 (AT2G29380), SnRK2.6 (AT4G33950), SnRK1a1 (AT3G01090) or UniProt database (https://www.uniprot.org/) under the following accession numbers: OsMODD (Q10Q07), TaAFP-B (B1B5D4), OsPP2C37 (Q7XP01) TaPP2C-a10 (A0A3B6ANL1). Residual data are available in the main text or the supplementary materials.

## Supplementary Materials

### Materials and Methods

Figs. S1 to S19

Tables S1 to S9

References (*62–88*)

## Notes

https://www.ebi.ac.uk/pride/

### Competing Interest Statement

The authors have declared no competing interest.

