## supplemental materials for "DOG1 prevents AFPs activation by AHG1 to control dormancy separately from ABA core signaling"

**The PDF file includes:**

Materials and Methods
Figs. S1 to S19
Tables S1 to S9
References (*62-88*)

#### Materials and Methods

##### Plant material, growth conditions, and seed batches productions.

Seeds of *Arabidopsis thaliana* (L.) Heynh were used in this study. The Near Isogenic Line NIL-DOG1, which contains the *DOG1* QTL from *Cape verde Island* (Cvi) in a *Landsberg erecta* (Ler) background, *dog1-1* (in NIL-DOG1 background), *dog1-2* (in Col-0 background) EMS null mutants, or the T-DNA insertion null single mutant *ahg1-5* (SALK_049885) *ahg3-2* (SALK_028132), as well as the double *ahg1-5 ahg3-2* or triple *ahg1-5 ahg3-2 dog1-2* were previously described (*13*, *20*, *26*). The *abi5-7* and *snrk2.2/3/6* mutant used as controls were previously described (*62*, *10*). The *afp1-3* (SALK_064533) *afp1-4* (SALK_098122C), *afp1-5* (SALK_020158C), *afp1-6* (SALK_005054) *afp2-1* (SALK_131676), *afp2-2* (SALK_145086C), *afp3-1* (SALK_037555), *afp3-2* (SALK_052114C), and *afp4-2* (SALK_208284C) are in the Col-0 background and were obtained from the NASC collection (Data source 1). Homozygous *afp* single mutants were isolated using allele-specific primers (Supplementary Table 3). The expression of *AFPs* full-length mRNA was investigated by RT-PCR, no corresponding transcript was amplified for *afp1-5*, *afp2-2* and *afp4-2*. The expression of *AFP1*, *AFP2* and, *AFP3* was reduced in *afp1-4*, *afp2-1* and *afp3-1* but not changed in *afp1-3/afp1-6,* and *afp3-2* compared to WT (fig. S15). Consequently, we considered *afp1-5*, *afp2-2*, and *afp4-2* as knock-out and *afp1-4* or *afp2-1* as knock-down alleles. Multiple mutants were obtained by crossing, and homozygous lines were selected from self-pollinated F2 progenies using gene specific primers for the T-DNA insertion mutant, and a CAPS marker for the *dog1-2* allele (Supplementary Tables 3, Supplementary Tables 4 and Data source 1). The hemizygous *35S::YFP:AFP2 (+/-)* line expressing full length AFP2 fused to YFP in its N-terminus was a gift from Prof. Ruth Finkelstein and corresponds to the previously described 4A2 line (*37*, *38*). Homozygous *YFP:AFP2* over-expressing seeds fail to complete maturation, are shrunken and non-viable impeding the physiological characterization of *35S::YFP:AFP2 (+/+)* progenies (*37*). Hemizygous *35S::YFP:AFP2 (+/-)* in *ahg1-5* or *ahg3-2* backgrounds were obtained by crossing, and homozygous *ahg* single mutants harbouring the *AFP2* transgene at the hemizygous state were selected from self-pollinated F2 progenies using alleles or transgene-specific primers (Supplementary Tables 3 and Supplementary Tables 4).

To produce seed batches for germination experiments, plants were cultivated on soil in a growth chamber with fluorescent light tubes with a 16 h day (150 µE $m^{-2}$ $s^{-1}$), 8 h night cycle at 50 % RH. Except when stated otherwise, growing temperature for plant lines in the Col-0 ecotype, was set at 21/18°C (day/night) until bolting and lowered to 16/14 °C during seed production and maturation to enhance seed dormancy at harvest (*63*). NIL-DOG1, *dog1-1* seed batches were matured at 21/18°C.

Seeds were harvested when approximately two third of the siliques turned brown by gently shaking the floral stem in paper bags to avoid inclusion of seeds from immature siliques. Biological replicates were defined as the progenies of a single mother plant. Non-seed contaminants were removed through sieving on a 355 µM mesh and non-filled (aborted/undeveloped) seed were mechanically sorted out by rolling seeds on paper several times. This treatment removed most of the black, wrinkled and non-viable seeds from *35S::YFP-AFP2* (+/-) progenies. Consequently, the proportion of transgenic genotypes in seed batches from *35S::YFP-AFP2* (+/-) progenies is expected to be around 66 % hemizygous and 33 % wildtype. Processed seed batches were equilibrated for 7 days in a container fixed at 21 °C and 50 % HR before the start of the experiments (WAH0) and, stored in these conditions during the full after-ripening kinetics.

**Germination assays**

For each genotype, approx. 50 seeds per biological replicate were sown on filter paper (Whatman grade 1) moistened with doubled distilled water in petri dishes. Seed plates were incubated under optimal conditions for Arabidopsis germination: in a transparent container fixed at 99 % RH which was placed inside a growth chamber with fluorescent light tubes with a 16 h day (80 µE $m^{-2}$ $s^{-1}$) / 8 h night cycle and a constant temperature of 22 °C. To follow dormancy alleviation through after-ripening, seed batches were sown at regular intervals along storage in fixed condition of 22 °C and 55 % RH. The full viability of seeds batches produced in our conditions for each genotype was verified by imbibing seeds in presence of 50 µM gibberellic acid ($GA_{4+7}$) followed by a stratification treatment before transfer to germination conditions (fig. S16). Fluridone and abscinazole-E3M were used to manipulate the endogenous ABA level at imbibition (*64*, *65*). Stock solutions of fluridone, abscinazole-E3M, and GA were dissolved in DMSO (final concentrations in the assays were 0.05 %). When imbibed on chemicals, the mock control corresponds to 0.05 % DMSO in water. Germination *per se* (radicle emergence) was scored after 7 days of incubation. Raw germination data are available in Data source 2.

**Transcript analysis**

Total RNA was extracted from 25 mg dry seeds using the RNAqueous™ Total RNA Isolation Kit (Invitrogen™ ThermoFisher Scientific, Germany) as previously described (*13*). Samples were treated with DNA-free™ kit (Invitrogen™ ThermoFisher Scientific, Germany) and 1 µg per RNA sample was used for reverse transcription using the FastGene® Scriptase II kit (Nippon Genetics EUROPE GmbH, Germany). Quantitative PCRs were performed using iQ SYBR® Green Supermix in a CFX Connect Real-Time PCR Detection System (Bio-Rad Laboratories, Inc., USA) as previously described (*66*). *PEX4* was used in all RT-qPCR and *TIP41* and *UBI10* in semi-quantitative PCR experiments as a validated internal housekeeping gene control for seed transcriptomic analysis (*41*). (Semi)-quantitative PCR Primers sequences are available in Supplementary Table 5. Raw data are available in Data source 3.

**Protein extractions**

Arabidopsis seeds or tobacco leaves samples were ground in liquid nitrogen with a mortar and pestle. Total proteins were solubilized in a buffer containing 50 mM HEPES pH 7.5; 2.5 % (w/v) SDS; 5 mM DTT under constant shaking at RT °C. Native protein complexes from tobacco leaves were solubilized in a buffer containing 50 mM Tris-HCl pH 7.9, 100 mM NaCl, 0.25 mM $MgCl_2$, 17.5 % glycerol, 1 mM ascorbate, 5 mM DTT, 2.5 % CHAPS, 2 mM PMSF, 20 µM MG132, 1 U $ml^{-1}$ macerozyme (Boehringer Ingelheim, Germany), 50 U $ml^{-1}$ DNase I (Roche, Switzerland), 1 % (v/v) of plant protease inhibitor cocktail (Sigma-Aldrich, Germany) under constant shaking in an ice-bath for 40 min. Protein extracts were clarified by repeated centrifugation until debris and oil free samples were obtained. Protein concentrations in the extracts were quantified using the

Pierce 660 nm Protein Assay Reagent kit (Thermo Fisher Scientific, Germany) against a BSA standard dilution curve and directly used for downstream applications.

**Protein gel and immuno-detection methods**

Total (10 to 30 µg) or purified (0.5 to 2 µg) proteins were separated by SDS-PAGE using home-made 10 to 12 % (w/v) acrylamide/bis-acrylamide matrix. For Western blots, proteins were transferred on nitrocellulose membranes (Protran 0.2 µM nitrocellulose; Amersham Biosciences, UK) using Bjerrum-Schaefer-Nielsen buffer (48 mM Tris, 39 mM glycine, 15 % methanol pH 9.2) via semidry transfer. After transfer, proteins were stained with Ponceau S before blocking with 3 % BSA for 1 h at RT°C. Primary and secondary antibody were incubated for 1 h at RT °C or overnight at 4 °C. Detection was performed using SuperSignal™ West Femto Maximum Sensitivity (Thermo ScientificTM) and a Chemidoc XP (Bio-Rad) or an Odyssey (LI-COR) imagining system. Primary, secondary antibodies and corresponding working dilutions used in this study are available in Supplementary Tables 6 and 7. Full scan images for every Western blot membrane; and protein gel presented in this study are available in Data source 4.

**Gene constructs**

Full-length coding DNA sequences (CDS) for *AFP1, AFP2, AFP3, AFP4, ABI1, HAB1, HAB2, HAI1, HAI2, HAI3, SnRK2.6* and *SnRK1a1* were amplified from cDNA of dry Col-0 seeds using gene-specific primers including the stop codon and cloned into pDONR201 or 207 using BP reaction (Invitrogen) as previously described (*26*). The pDONORs entry clones for *AHG1*, *AHG3* and *ABI2* were previously described (*26*). The $AHG1_{D123\_149A}$ inactive variant was obtained using QuikChange II XL Site-Directed Mutagenesis Kit (Agilent). Primer sequences are available in Supplementary Table 8. Correct sequences were verified by Sanger sequencing at Microsynth SeqLab. pDONR entry clones were recombined into destination vectors using LR reaction (Invitrogen, USA).

**Generation of *35S:YFP:SnRK2.6* transgenic lines**

The CDS (including stop codon) of *SnRK2.6* was recombined from entry clones in the pEarlyGate104 vector (*67*) and transfected into *Agrobacterium tumefaciens* GV3101 pMP90 using electroporation before transformation of NIL-DOG1 and *dog1-1* using floral-dipping method as described in (*66*). Homozygous single insertion lines were selected at the T3 generation through their resistance to phosphinothricin and genotyping by PCR. Two independent lines in each background were isolated (lines 1 and 2 are in NIL-DOG1 and lines 3 and 4 are in *dog1-1*). Further comparative assays used line 2 and 4 based on a similar accumulation of the fusion protein in dry seed.

**Dissection of *A. thaliana* embryos and localization analysis.**

Six or 24 hours imbibed seeds in dd$H_2O$ were transferred onto a glass plates and intact embryos were isolated manually from surrounding tissue under a binocular as described in (*68*). Embryos were transferred onto microscope glass slides in water, for localization analyses using a Leica SP8 confocal microscope (Leica Microsystems, Mannheim, Germany) via a 63x water objective. YFP was excited at 514 nm with an argon laser, and emission was detected between 525-545 nm using a HyD detector. Imaging was conducted using following settings: format 1024 x 1024, bidirectional scan 400 Hz, pinhole of 1 airy unit (AU), digital zoom 1-3.

**SnRK2.6 purification from dry seed and kinase assays.**

Protein were extracted from 50 mg dry seed material as stated above with the following modification: protein were suspended in 1 mL of ice-cold extraction buffer (50 mM HEPES pH 7.5; 150 mM NaCl, 10 µM ATP, 1 % (w/v) Triton X-100, 5 mM DTT, 1 mM EGTA; 1 mM EDTA; 50 U/ml DNAse I, 1 U/ml Macerozyme, 20 mM β-glycerophosphate, 50 mM NaF, 1 mM $Na_3VO_4$, 1 % plant protease inhibitor cocktail, 20 µM MG132, 50 µM ABA), sample were vortexed, and incubated for 10 min at RT under gentle shaking before clarification by centrifugation. After quantification, total protein amount was normalized to 0.75 mg/mL. For each assay, 25 µl of agarose beads coupled to a GFP antibody (GFP-Trap®, ChromoTek) corresponding to 10 µg GFP-binding capacity was equilibrated 3 times in 50 mM HEPES pH 7.5; 150 mM NaCl, 5 mM DTT, 1 mM EGTA; 1 mM EDTA. One milligram of total protein was mix with beads followed by incubation under constant rotation for 2 h at 4 °C. After centrifugation (200 x g for 1 min) beads were washed twice with 500 µl of equilibration buffer and a last wash was performed using the kinase buffer (50 mM HEPES pH 7.5; 1 mM EGTA; 10 mM MgCl2; 1 mM DTT; 20 mM β-glycerophosphate). After the last centrifugation step, the supernatant was discarded, and the beads were resuspended in 50 µl (final volume) of kinase buffer. A 15µl aliquot was mixed with 15 µl of 2x Laemmli buffer (4x stock: 1 % SDS, 10 % glycerol, 0.005 % bromophenol blue, 62.5 mM Tris-HCl, pH 6.8, 355 mM 2-mercaptoethanol) and used for Western blot analyses. The residual 35 µl were mixed with 15 µl kinase buffer and the reaction was started by addition of 1.5 µl NaCl (150 mM), MBP 1.5 µl (10 mg/ml), 0.15 µl ATP (10 mM), 2 µCi $^{32}P$ γ-ATP. After 1 hour incubation at RT°C, the reaction was stopped by addition of 15 µl of 4x Laemmli buffer before SDS-PAGE separation. Gels were stained and fixed using Coomassie staining solution and placed in an exposure cassette with a phosphor screen for 1 hour. The incorporation of $^{32}P$ into the MBP substrate was analyzed using a Typhoon™ FLA700 Scanner (GE Healthcare). The images were taken using the Phosphorimaging mode, at a pixel size of 25 µM and a latitude of 4. The voltage, which was applied to the photomultiplier tube was set at 750 V. Full scan images of autoradiogram and total staining are available in Data source 4.

**Yeast two-hybrid assay**

The CDS (including stop codon) of *AFP1*, *AFP2*, *AFP3*, *AFP4*, *SnRK2.6* and, *ABI1*, *HAB1*, *HAB2*, *HAI1*, *HAI2*, *HAI3* were recombined from entry clones in the pAS2-gateway (GAL4 BD fusion) and pACT2-gateway (GAL4 AD fusion) vectors (modified from Clontech) respectively. pACT2:AHG1 pACT2:AHG3 and pACT2:ABI2 were previously described (*26*). Interaction assays were performed in the yeast strain PJ69-4alpha as previously described (*26*). Interactions were monitored *via* drop tests on -LWH medium (-LWH) lacking L, W and His (H) with 5 mM and 10 mM 3-aminotriazole (3-AT) in dilutions of $10^{-1}$, $10^{-2}$ and $10^{-3}$. Yeast was grown at 30 °C and photos were recorded after 2 or 4 days for -LW or -LWH plates, respectively.

**Tobacco leaves transient transformation**

Electro-competent *Agrobacterium tumefaciens* GV3101 (pMP90) cells were transformed with binary destination vectors and grown at 28 °C (*69*). Overnight cultures inoculated with single colonies were cultivated under constant shaking at 28 °C. Bacteria were diluted to a final $OD_{600}$ of 0.5 in activation medium for tobacco infiltration (10 mM $MgCl_2$, 10 mM MES pH 5.7, 100 µM acetosyringone) and mix with the p19 helper strain at an $OD_{600}$ of 0.4 (*70*). Four weeks old *Nicotiana benthamiana* leaves were infiltrated with a syringe.

**Co-immunoprecipitation (CoIP)**

The full-length CDS (including stop) of *AFP1, AFP2, AFP3, AFP4, SnRK2.6* and *SnRK1a1* were recombined from entry clones in the pBat-TL-Venus-GW (gift from Dr. Philip Känel, Fraunhofer IME, Münster, Germany) vector while *AHG1, AHG3* and *ABI2* were recombined in pAlligator2 vector (*71*). Recombined vectors were used for tobacco co-infiltration and native protein complexes were purified from 48 h infiltrated leaves. For each pull-down, 25 µl of agarose beads coupled to a GFP antibody (GFP-Trap®, ChromoTek) was equilibrated in a buffer containing 50 mM Tris-HCl pH 7.9, 100 mM NaCl, 0.25 mM $MgCl_2$, 17.5 % glycerol, 1 mM ascorbate, 5 mM DTT. Equilibrated beads were dispensed in 2 ml of native extract containing 3 mg of protein and incubated under rotation at 4 °C for 45 min. After incubation, beads were separated from non-bound fractions by centrifugation, and washed three times with 500 µl of equilibration buffer. Bound protein complexes were eluted from the beads by adding of 4x Laemmli buffer, followed by 5 min incubation at RT °C. Samples were diluted 4 times with $ddH_2O$ and boiled at 95 °C before SDS-PAGE separation. For each interaction paring, input and elution fraction were analysed on a single membrane. Venus (Bait) fusion proteins were detected using Anti-GFP antibody. After stripping of membranes, 3xHA (Prey) fusion proteins were detected on the same membrane using Anti-HA antibody. The Ponceau S staining was used as loading control to control the quality of the Venus-tagged fusion protein enrichment (ratio signal Anti-GFP/ RbcL).

**Production and purification of recombinant protein**

Preparation of recombinant 6xHIS:MBP:AHG1 and 6xHIS:MBP:$AHG1_{D123\_149A}$ was performed as previously described (*26*). The purity and the specific activity towards of the enzymes used in this study was followed by SDS-PAGE and using the RRA{pT}VA standard peptide respectively (fig. S12A).

***In vitro* phosphatase assays**

The RRA{pT}VA standard peptide was obtained from Promega (ref. V248A). Synthetic phosphorylated peptides of 14 amino acids from AFP1 modified at S115 (GLMRTT{pS}LPAESEE), AFP2 modified at S112 (GLERTT{pS}LPAEMEE), and SnRK2IIIs modified at a conserved serine corresponding to S171 in SnRK2.6 (KSSVLH{pS}QPKSTVG) were obtained from Biomatik, LLC (USA). Reactions were performed in flat-bottom 96 wells plate in a final volume of 50 µL. The phosphatase reactions were started by addition of 4.4 µM recombinant 6xHIS:MBP:AHG1 or 6xHIS:MBP:$AHG1_{D123\_149A}$ proteins in the reaction mix containing 50 mM HEPES buffer pH 7.5; 10 mM $MgCl_2$ and the indicated concentration of the corresponding substrate peptide. Reactions were performed for 5 to 30 min at RT and stopped by addition of 100 µl of BIOMOL® Green (Enzo Life Sciences, Inc., Germany). The dye was developed for 30 min at room temperature before measuring the absorbance at 630 nm in a Tecan Infinite M200 Pro plate reader (Tecan Group, Switzerland). Specific activities were calculated using a serial dilution of free phosphate as standard. As positive control for the dephosphorylation of the SnRK2IIIs peptide, a reaction was performed using 0.07 µM of commercial Calf Intestinal Alkaline Phosphatase (Thermo Fisher Scientific, Germany). Raw activity data are available in Data source 5.

**Proteolytic digestion**

Total protein extracts form NIL-DOG1 or *dog1-1* dry or 6 h imbibed seeds were digested to peptides using the Filter Aid Sample Preparation (FASP) method (*72, 73*). Proteins (1 mg) were diluted in 8 M urea in 0.1 M Tris HCl pH 8.5 (to a final volume 1 ml) and reduced by addition of DTT at a final concentration of 10 mM for 1 h at RT °C. Cysteines were alkylated by addition of CAA at a final concentration of 45 mM and incubated for 1 h in the dark at RT °C. The excess of CAA was quenched by addition of DTT at a final concentration of 83 mM for 1 h in the dark at RT °C. The whole mixture was transferred on Amicon 10 kDa centrifugation filter (Millipore, Darmstadt, Germany) and diluted 3 times with 50 mM Ammonium bicarbonate (ABC) buffer. The volume was reduced to 500 µL by centrifugation at 3,000 x g and further diluted 6 times with ABC buffer. The volume was further reduced to 250 µl by centrifugation and proteins were digested by addition of 2 µg trypsin (T6567, Sigma-Aldrich, Darmstadt, Germany) and incubation at 37 °C ON under agitation. Digested peptides were recovered in the flow-through by centrifugation. Concentrator filters were washed twice with 0.5 M NaCl followed by centrifugation. The flow through from wash fractions was combined with recovered peptides. For the targeted analysis of YFP:SnRK2.6 phosphorylation status, the sample were prepared as for the kinase assays with the following modification: the β-glycerophosphate was omitted in extraction buffer and the last bead wash was performed with equilibration buffer. Elution was performed by incubating beads in 0.1 % of TFA for 5 min before supernatant harvest and neutralization with an equal volume of UA buffer (8 M urea in 0.1 M Tris–HCl pH 8.5). Eluted proteins were reduced for 30 min in the dark by adding DTT at a final concentration of 12 mM was and subsequently alkylated for 1 h in the dark by adding chloroacetamide at a final concentration of 44 mM. The excess of CAA was quenched for 30 min in the dark by adding DTT at a final concentration of 120 mM. The mixture was diluted seven times with ABC buffer, before addition of 1 µg trypsin and overnight incubation at 37 °C.

**Mass spectrometry sample preparation**

Peptides from total protein extracts were acidified by addition of formic acid at a final concentration of 0.5 % and desalted using Sep-Pak SPE 1 cc/100 mg (Waters, Eschborn, Germany) as previously described (*74*). Ten micrograms of desalted peptides were processed for total proteome analysis by pre-fractionation in three fractions using Empore Styrenedivenylbenzene Reversed Phase Sulfonate material (SDB-RPS; 3 M) as previously described (*75*). Phosphopeptide enrichment from total protein extracts was performed by solving 400 µg of desalted peptides in 50 % Acetonitrile (ACN), 1 % trifluoroacetic acid (TFA) supplemented with 20 mg/mL of 2,5-Dihydroxybenzoic acid (DHB) and subjected to titanium dioxide ($TiO_2$) affinity chromatography using 6 mg of $TiO_2$ beads (5 mm; GL Sciences) per samples as previously described (*74*). Elution from $TiO_2$ or agarose coupled to a GFP antibody beads were further desalted with StageTips (Empore C18; 3M) as previously described (*76*). After desalting, peptide samples were dried in a centrifugal evaporator.

**LC-MS/MS Data acquisition**

Before measurement, peptides were resuspended in A* buffer (2 % ACN, 0.1 % TFA). Proteomic samples were measured by liquid chromatography-tandem mass spectrometry (LC-MS/MS) using Data Dependant Acquisition (DDA) on an EASY-nLC 1000 system (Thermo Fisher Scientific, Germany) coupled to an Orbitrap Q Exactive Plus mass spectrometer (Thermo Fisher Scientific, Germany) as previously described (*74*). For targeted analysis of YFP:SnRK2.6 phosphorylation status the samples were split in two and measured on an EASY-nLC 1200 system (Thermo Fisher Scientific, Germany) coupled to an Orbitrap Exploris 480 mass spectrometer (Thermo Fisher Scientific, Germany) as previously described for DDA (*77*) and using the Parallel Reaction Monitoring (PRM) mode targeting peptides summarized in Supplementary Table 9.

**MS data processing and analysis**

Raw LC-MS/MS spectra were processed using MaxQuant (version 1.6.17.0) (http://www.maxquant.org/) against the Arabidopsis Araport11 database (*78)* with label-free quantification (LFQ) enabled (*79*). Common protein contaminant and decoy sequences were automatically added during the search. Trypsin cleavage specificity was required with a maximum of two missed cleavages allowed, and a minimal peptide length of seven amino acids. Carbamidomethylation of cysteine residues was set as fixed, and oxidation of Met and protein N-terminal acetylation as variable modifications. For the analysis of phosphopeptide enriched samples, phosphorylation of Ser, Thr, and Tyr were added as additional variable post-translational modifications. False discovery rate cut-off was set at 1 %. A total of 5004 Arabidopsis protein groups and 4848 unique p-peptides (FDR < 1 %, score > 40 and delta score > 6) covering 1616 phosphorylation sites were quantified. MaxQuant output files were further analysed using version 1.6.6.0 of Perseus (*80*). For both proteome and phosphoproteome measurements, the reproducibility between replicates for a given genotype and condition was very high as demonstrated by Pearson correlation coefficients calculated in Perseus (fig. S17-18). Protein LFQ intensities (total proteome analysis) or peptides intensities (phospho-proteome analysis) were $log_2$ transformed, and the data was filtered for valid values in at least two out the three replicates of one genotype in either dry or imbibed conditions. After filtering, 4324 protein groups and 984 phosphopeptides (for which the exact phosphorylated residues could be determined on 828 sites) were retained for statistical analysis. MS outcome metrics are available as Supplementary Table 1. Annotation for protein localization were performed in Perseus using the SUBA4 consensus database. Missing values were imputed from normal distribution using a width of ($log_2$) 0.5 or 0.6 and, a downshift of ($log_2$) 2 or 1.95 for total proteomes or phosphoproteome datasets respectively (fig. S19). We used an empirical Bayes method to identify significant changes in protein or p-peptides abundance between genotype in dry or 6 h imbibed seeds. Fold changes and *P*-values were calculated using the LIMMA ("Linear Models for Microarray Data") package on R (*81*). Data analysis outcomes are available as Data source 6. Gene Ontology were performed in

Cytoscape using a Hypergeometric test with a significance level of 0.05 after correction by a Benjamin & Hochberg FDR test in the BiNGO app (*82, 83*). A seed dry or imbibed proteome obtained from the Arabidopsis Draft Proteome was used as reference as background set for GO analysis (*84).* Protein annotated as implied in ABA responses were retrieved using the GO:009737 terms in Amigo2 database (https://amigo.geneontology.org/amigo). For targeted analysis of YFP:SnRK2.6 phosphorylation status, raw LC-MS/MS spectra from both acquisition methods (DDA and PRM) were processed together as separate parameter groups using the MaxQuant with the settings described above expected that LFQ quantification was disabled for PRM data. Localization probabilities were further manually filtered retaining entries with a score superior to 60 to ensure high confidence identification and localization. Data analysis outcomes are available as Data source 7.

**Protein sequence, phylogeny, and structures analyses**

Protein amino acid sequence alignments and phylogenetic trees were performed in Mega11 using the ClustalOmega and the maximum likehood joining algorithms. Structures of Arabidopsis SnRK2.6 in complex with HAB1 were obtained from the PDB repository (https://www.rcsb.org/) with the accession number 3UJG and modelling of AHG1 was obtained from the Alphafold repository (https://alphafold.ebi.ac.uk/). Superimposition of protein structures was performed using PDBeFold (https://www.ebi.ac.uk/msd-srv/ssm/). Visualization and analyses were performed using PyMOL software (DeLano Scientific LLC).

**Image and statistical analysis**

Western blot, autoradiogram and gel signal intensities were obtained by image densitometry analysis using ImageJ. Values and calculations are available in Data source 8. Differences in paired or unpaired comparison were tested using a student t-test. For multiple comparisons, germination percentage values were arcsin square root transformed to accomplish the prerequisites of normality before applying a one-way ANOVA with a Tukey Honestly Significance Difference (HSD) post-hoc test. Tests were performed in the Origin2023 software. Bar graphs were built in GraphPrism V10.2.1.

**Accession numbers and data availability**

Mass spectrometry proteomics data have been deposited in the ProteomeXchange Consortium (http://proteomecentral.proteomexchange.org) via the JPOSTDB partner repository with the data set identifier PXD046985 and PXD053457. Sequence data from this article can be found in The Arabidopsis Information Resource-TAIR database (www.arabidopsis.org) under the following accession numbers: AFP1 (AT1G69260), AFP2 (AT1G13740), AFP3 (AT3G29575), AFP4 (AT3G02140), AHG1 (AT5G51760), AHG3 (AT3G11410), ABI1 (AT4G26080), ABI2 (AT5G57050), HAB1 (AT1G72770), HAB2 (AT1G17550), HAI1 (AT5G59220), HAI2 (AT1G07430), HAI3 (AT2G29380), SnRK2.6 (AT4G33950), SnRK1a1 (AT3G01090) or UniProt database (https://www.uniprot.org/) under the following accession numbers: OsMODD (Q10Q07), TaAFP-B (B1B5D4), OsPP2C37 (Q7XP01), TaPP2C-a10 (A0A3B6ANL1).

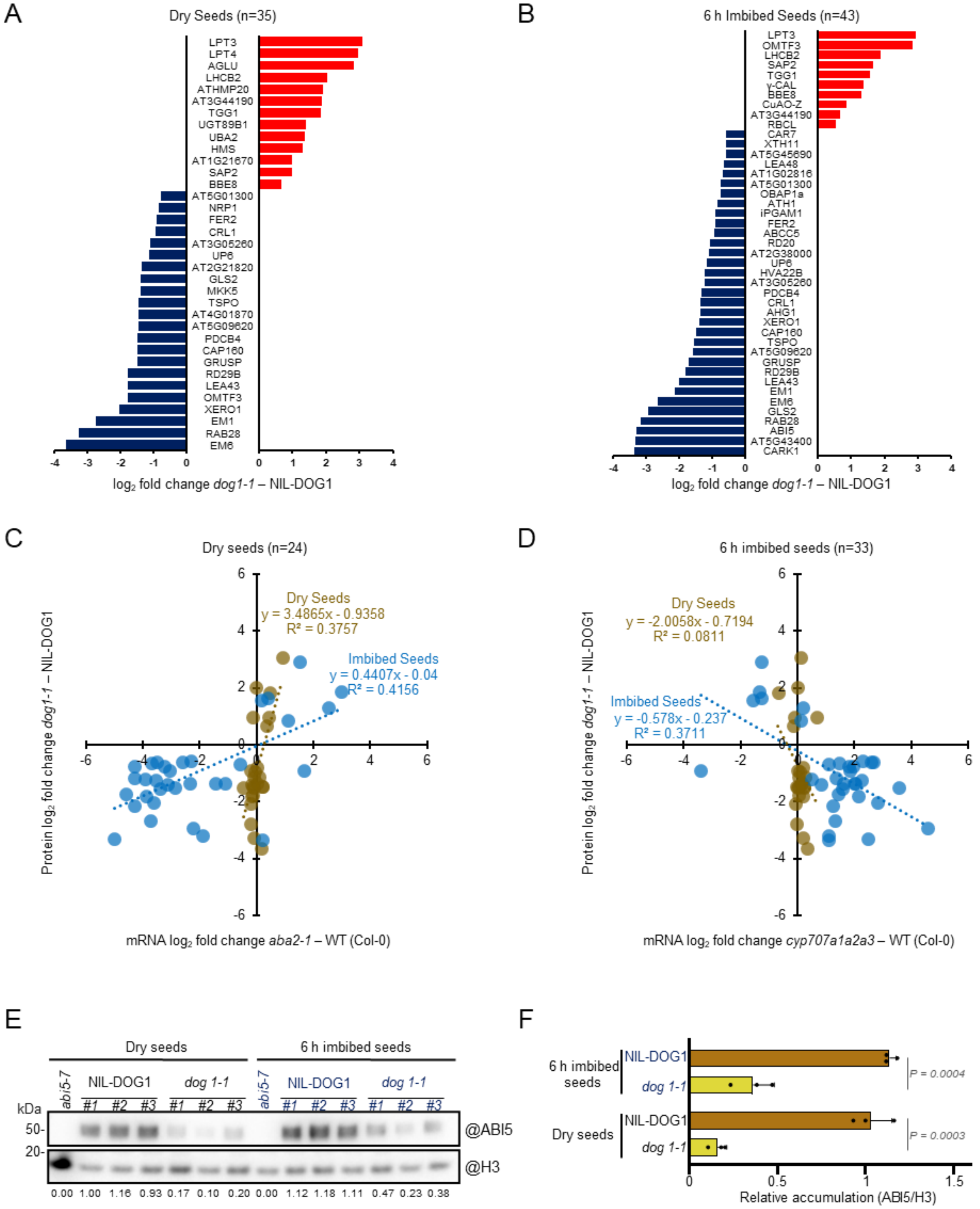
A
Dry Seeds (n=35)
LPT3
LPT4
AGLU
LHCB2
ATHMP20
AT3G44190
TGG1
UGT89B1
UBA2
HMS
AT1G21670
SAP2
BBE8
AT5G01300
NRP1
FER2
CRL1
AT3G05260
UP6
AT2G21820
GLS2
MKK5
TSPO
AT4G01870
AT5G09620
PDCB4
CAP160
GRUSP
RD29B
LEA43
OMTF3
XERO1
EM1
RAB28
EM6
log2 fold change dog1-1 – NIL-DOG1
B
6 h Imbibed Seeds (n=43)
LPT3
OMTF3
LHCB2
SAP2
TGG1
γ-CAL
BBE8
CuAO-Z
AT3G44190
RBCL
CAR7
XTH11
AT5G45690
LEA48
AT1G02816
AT5G01300
OBAP1a
ATH1
iPGAM1
FER2
ABCC5
RD20
AT2G38000
UP6
HVA22B
AT3G05260
PDCB4
CRL1
AHG1
XERO1
CAP160
TSPO
AT5G09620
GRUSP
RD29B
LEA43
EM1
EM6
GLS2
RAB28
ABI5
AT5G43400
CARK1
log2 fold change dog1-1 – NIL-DOG1
C
Dry seeds (n=24)
Dry Seeds
y = 3.4865x - 0.9358
R² = 0.3757
Imbibed Seeds
y = 0.4407x - 0.04
R² = 0.4156
Protein log2 fold change dog1-1 – NIL-DOG1
mRNA log2 fold change aba2-1 – WT (Col-0)
D
6 h imbibed seeds (n=33)
Dry Seeds
y = -2.0058x - 0.7194
R² = 0.0811
Imbibed Seeds
y = -0.578x - 0.237
R² = 0.3711
Protein log2 fold change dog1-1 – NIL-DOG1
mRNA log2 fold change cyp707a1a2a3 – WT (Col-0)
E
Dry seeds
6 h imbibed seeds
abi5-7
NIL-DOG1
dog 1-1
#1 #2 #3 #1 #2 #3
kDa
50-
20-
@ABI5
@H3
0.00 1.00 1.16 0.93 0.17 0.10 0.20 0.00 1.12 1.18 1.11 0.47 0.23 0.38
F
6 h imbibed seeds
Dry seeds
NIL-DOG1
dog 1-1
P = 0.0004
P = 0.0003
Relative accumulation (ABI5/H3)

**Fig. S1: Typical ABA responses are altered in *dog1-1* dry and early imbibed seeds.** (**A**&**B**) Differential accumulation of significantly regulated proteins implied in ABA responses between genotypes in dry (**A**) or imbibed (**B**) seeds (protein abundances are expressed as $log_2$ fold changes, n= 3 biological replicates, *P*-value < 0.05). Up- and downregulated proteins in the *dog1-1* mutant are indicated in red and blue, respectively. (**C&D**) Comparison of regulation observed at the proteome level in *dog1-1* seeds with the regulation at the transcriptome level in the ABA deficient *aba2-1* mutant (**C**) or in the ABA over-accumulating triple *cyp707a1a2a3* mutant (**D**). The dashes lines show linear regression in dry (brown) or imbibed (blue) seeds labelled with their correlation coefficient (r) and trajectory equation. Transcriptomic data was obtained from the study by Okamoto et al. (*30*). (**E&F**) Quantification of ABI5 by Western blot analysis in total protein extracts from dry or 6 hours imbibed NIL-DOG1 and *dog1-1* seeds. The # indicates independent replicates. The bar chart shows protein accumulations normalized to the H3 loading control (Means±SD, n=3 biological replicates, unpaired t-test *P*-value is indicated) and presented as relative values to NIL-DOG1 levels (arbitrary mean value of 1 for replicate #1 dry seed).

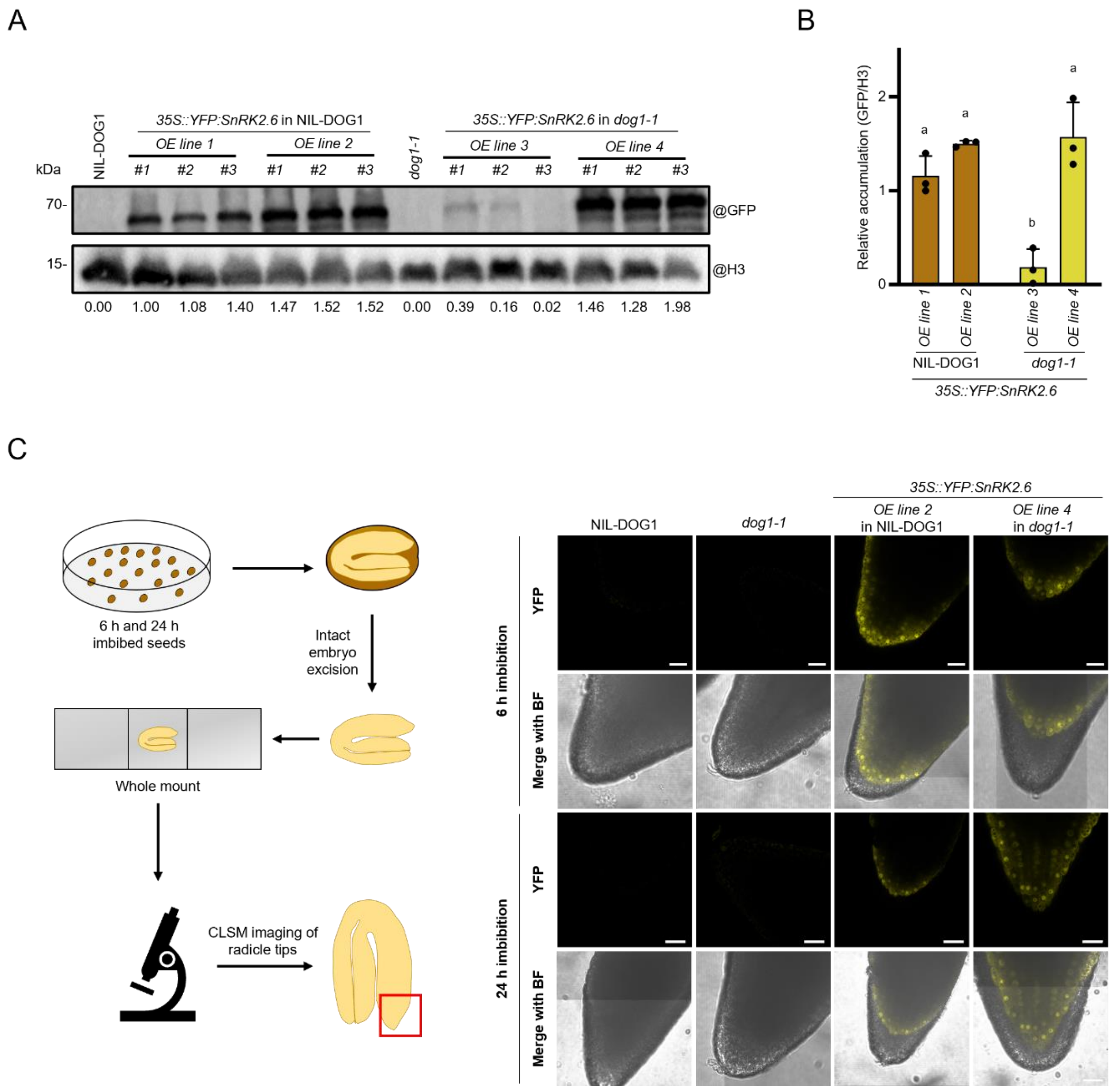


**Fig. S2: Analyses of *YFP:SnRK2.6* overexpression lines.** (**A**) Quantification of YFP:SnRK2.6 by anti-GFP Western blot analysis in total protein extracts from dry seeds of NIL-DOG1 and *dog1-1* overexpression lines. Non-transgenic NIL-DOG1 and *dog1-1* served as specificity controls for the antibody. The # indicates independent replicates. (**B**) The bar chart shows protein accumulations of YFP:SnRK2.6 normalized to the H3 loading control (Means±SD, n=3 biological replicates). Data are presented as relative values to NIL-DOG1 level (arbitrary mean value of 1 for replicate #1). Letters on the top of the bars indicate significantly different groups using a one-way ANOVA with a Tukey HSD test (α=0.05). (**C**) Confocal Laser Scanning Microscopy (CLSM) analysis of YFP:SnRK2.6 localization in intact embryos excised after 6 and 24 h imbibition. Shown are the YFP emission channel and a merged image with the bright field channel (BF). Scale bars represent 20 µm.

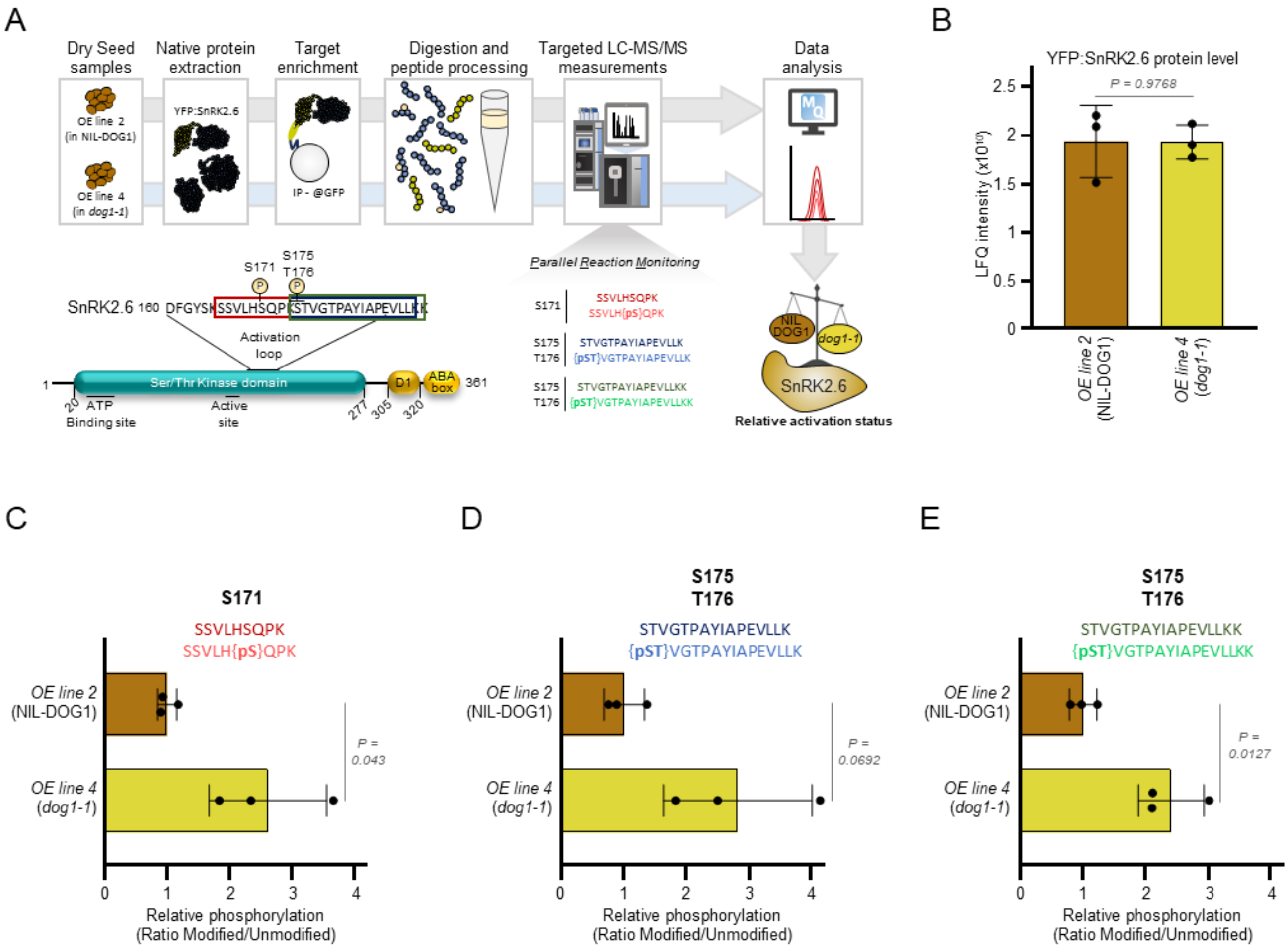


**Fig. S3: Targeted analysis of YFP:SnRK2.6 phosphorylation in NIL-DOG1 and *dog1-1* seeds.** (**A**) Workflow of the Parallel Reaction Monitoring (PRM) experiment. YFP:SnRK2.6 was enriched from transgenic NIL-DOG1 and *dog1-1* dry seed extracts. Phosphorylated and unmodified tryptic peptides from the class III SnRK2 activation loop (indicated in red, blue and green) covering the residues S171 and S175/T176 were targeted during measurement. Note that the green peptide is a miss-cleaved peptide species related to the blue one (**B**) Bar chart showing the loading of the sample based on protein LFQ intensity of SnRK2.6 purified from NIL-DOG1 and *dog1-1* background (loading control). (**C-E**) Bar charts showing the phosphorylation status of S171 (**C**), and S175/T176 (**D&E**) calculated through the ratio of modified/unmodified peptide intensities (Means±SD, n=3 biological replicates, unpaired t-test *P*-values are indicated) and presented as relative values to NIL-DOG1 (arbitrary mean value of 1).

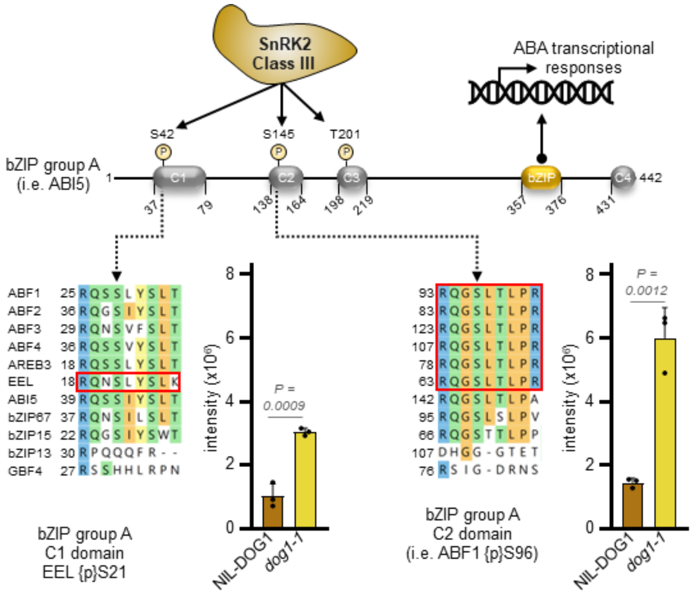


**Fig. S4: Conserved phosphorylated peptides of clade A bZIPs are more abundant in *dog1-1* seeds.** Schematic representation of group A bZIP proteins (using ABI5 as an example). The tryptic p-peptides identified in this study arose from the conserved C1 and C2 domains (indicated by the dashed arrows) and are highlighted in red within the sequence alignment sections. The peptide from group A bZIPs' C1 domain is unique to EEL but the peptide from the C2 domain can derive from any of the ABFs, AREB3 or EEL. Activation of ABA transcriptional responses by group A bZIPs is promoted by SnRK2IIIs mediated phosphorylation of conserved serine residues (i.e. S42, S145 and T201 in ABI5) within C1-3 domains (*35*). The bar charts on the left of the alignments shows MS intensities of the corresponding p-peptides in NIL-DOG1 and *dog1-1* dry seeds (Means±SD, n=3 biological replicates, unpaired t-test *P*-values are indicated).

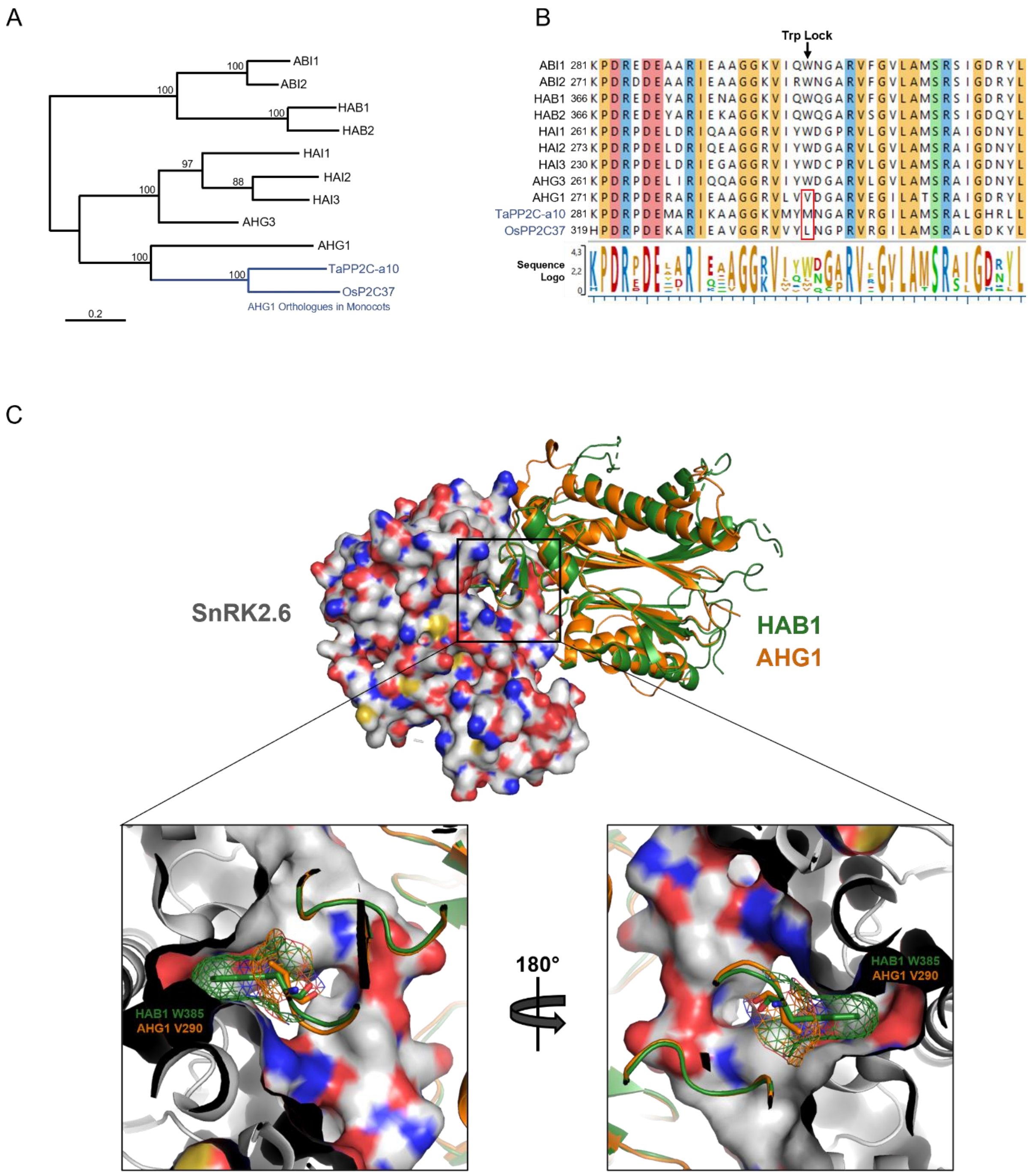

A
ABI1
ABI2
HAB1
HAB2
HAI1
HAI2
HAI3
AHG3
AHG1
TaPP2C-a10
OsP2C37
AHG1 Orthologues in Monocots
0.2
B
Trp Lock
ABI1 281 KPDREDEAARIEAAGGKVIQWNGARVFGVLAMSRSIGDRYL
ABI2 271 KPDRDDEAARIEAAGGKVIRWNGARVFGVLAMSRSIGDRYL
HAB1 366 KPDREDEYARIENAGGKVIQWQGARVFGVLAMSRSIGDRYL
HAB2 366 KPDREDEYARIEKAGGKVIQWQGARVSGVLAMSRSIGDQYL
HAI1 261 KPDRPDELDRIQAAGGRVIYWDGPRVLGVLAMSRAIGDNYL
HAI2 273 KPDRPDELDRIQEAGGRVIYWDGARVLGVLAMSRAIGDNYL
HAI3 230 KPDRPDELDRIEGAGGRVIYWDCPRVLGVLAMSRAIGDNYL
AHG3 261 KPDRPDELIRIQQAGGRVIYWDGARVLGVLAMSRAIGDNYL
AHG1 271 KPDRPDERARIEAAGGRVLVVDGARVEGILATSRAIGDRYL
TaPP2C-a10 281 KPDRPDEMARIKAAGGKVMYMNGARVRGILAMSRALGHRLL
OsPP2C37 319 HPDRPDEKARIEAVGGRVVYLNGPRVRGILAMSRALGDKYL
Sequence Logo
C
SnRK2.6
HAB1
AHG1
HAB1 W385
AHG1 V290
180°

**Fig. S5: Absence of the PP2CAs' conserved Trp-lock residue in AHG1 and its monocotyledon orthologues precludes hindrance of SnRK2s catalytic cleft.** (**A**) Phylogenetic tree of the 9 PP2CAs from Arabidopsis, and AHG1 orthologues from wheat (TaPP2C-a10), and rice (OsPP2C37). (**B**) Section of amino acid sequence alignment of Arabidopsis PP2CAs with TaPP2C-a10 and OsPP2C37. The position of the conserved Trp-lock residue is marked by an arrow and the amino acid substitution in AHG1 and AHG1-like is indicated in red. (**C**) Three-dimensional representation of the inhibition of SnRK2IIIs by PP2CAs through hindrance of the catalytic cleft by the Trp-lock residue. Structural modelling of AHG1 is superimposed to HAB1 subunit in complex with SnRK2.6. The bottom panels show an enlarged view of the squared region focusing on SnRK2.6 catalytic cleft occupancy by PP2CAs Trp-lock residue. The surface of SnRK2.6 is shown and HAB1 or AHG1 (AA 138-416) appears as green or orange ribbons respectively. HAB1 W385 and AHG1 V290 are shown as sticks. Meshes in the bottom panels represent the electron density map.

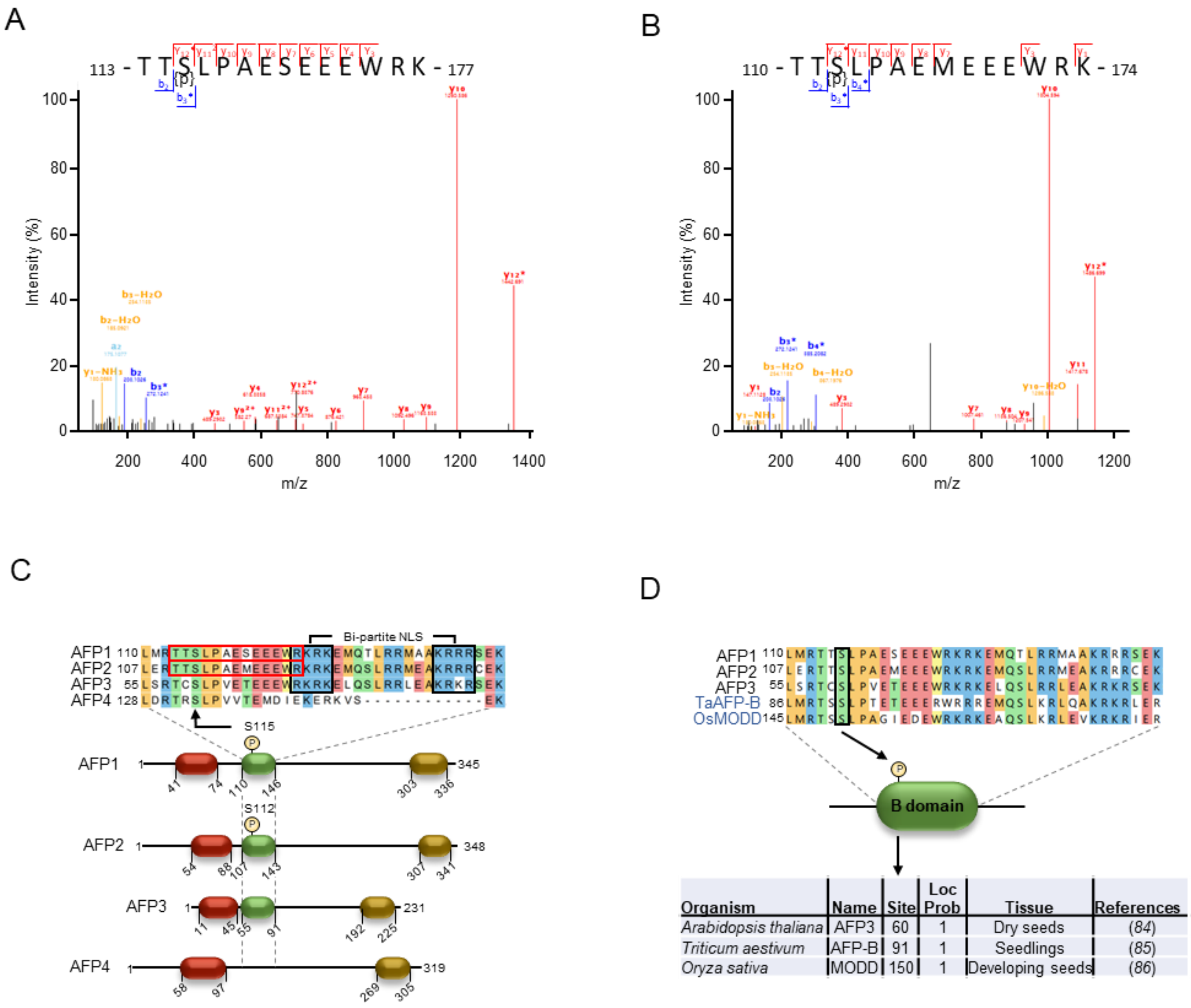


**Fig. S6: Phosphorylation of a serine of AFPs B domain is conserved in monocotyledon AFP orthologues.** (**A-B**) Annotated $MS^2$ spectra of the tryptic peptides phosphorylated at AFP1 S115 (**A**) and AFP2 S112 (**B**) with the identified fragment ions and respective start and end positions of the peptides as well as modified residues within AFP1 and 2 indicated additionally in the inset. b ions blue, y ions red, fragment ions with losses yellow. (**C**) Schematic representation of Arabidopsis AFP1-4 proteins and amino acid sequence alignment of the conserved B-domain. The tryptic p-peptides, which are less abundant in *dog1-1* compared to NIL-DOG1, are highlighted in red and have AFP1 or AFP2 as unique precursors. The modified conserved residue is linked to the alignment by an arrow. NLS = Nuclear Localization Sequence (black boxes). (**D**) Section of amino acid sequence alignment of Arabidopsis AFP1-3, and monocotyledon AFPs orthologues from wheat (TaAFP-B), and rice (OsMODD). The modified serine in the B-domain (box and arrow) is conserved in TaAFP-B and OsMODD. This residue was shown to be modified by phosphorylation in previous phosphoproteomic studies in AFP3 and monocotyledon AFP orthologues (*84-86)*.

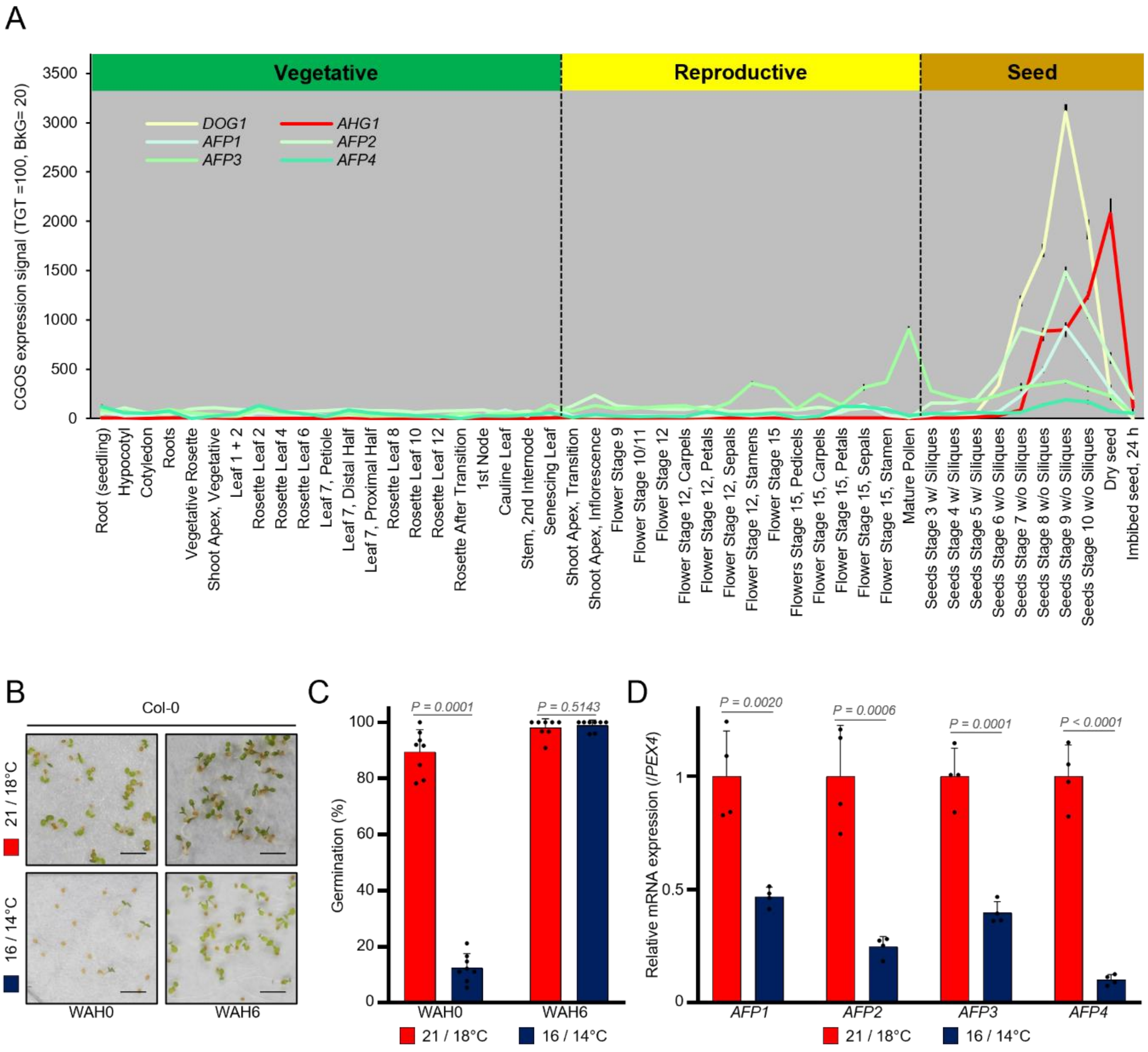


**Fig S7: *AFPs* expression is developmentally and environmentally regulated.** (**A**) Relative mRNA accumulation of *AFPs* during diverse developmental stages of Arabidopsis. Shown are normalized relative expression levels using the GeneChip Operating Software (CGOS) methods with a global scaling to target signal (TGT) = 100, and a background (BkG) of 20. The graphs are based on publicly available data from the eFP browser (http://bar.utoronto.ca/efp/cgi-bin/efpWeb.cgi). (**B**) Representative pictures (scale bars represent 2 mm) and (**C**) germination capacity of Col-0 seeds cultivated at 16/14 or 21/18 °C after flowering at 0 or 6 weeks after harvest (WAH), (Means±SD, n=8 biological replicates, unpaired t-test *P*-value is indicated). (**D**) Expression of *AFPs* transcripts in freshly harvested dry Col-0 seeds cultivated at both temperature regimes. The relative expression was normalised to the level of seed maturated at 21/18 °C (arbitrary value of 1, dashed line) for each *AFP* (Means±SD, n=4 biological replicates, unpaired t-test *P*-values are indicated).

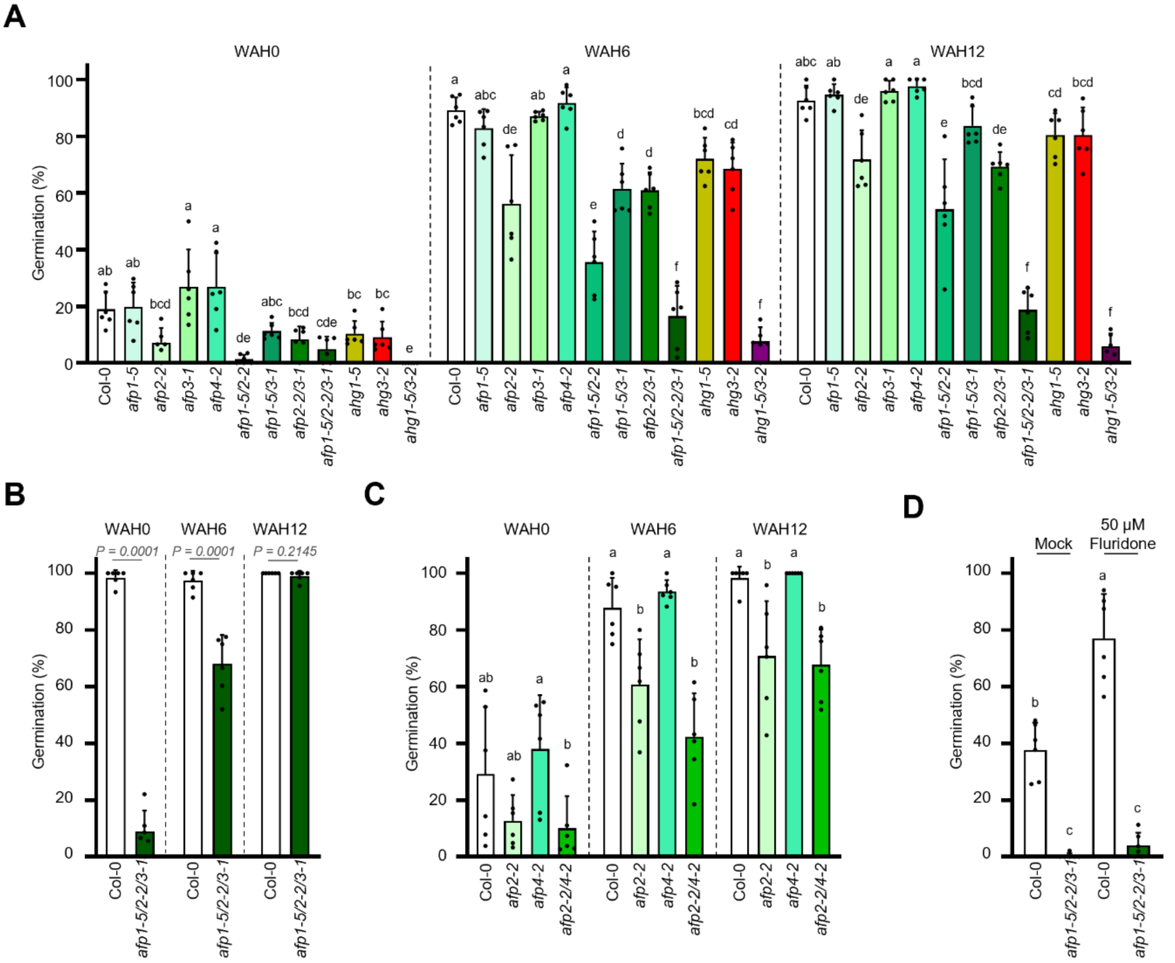


**Fig S8: AFPs are partially redundant regulators of seed dormancy in Arabidopsis. (A)** Germination capacity of single and multiple *afp* mutants at harvest, and during after-ripening at 6 and 12 weeks after harvest (WAH) (Means±SD, n=6 biological replicates). (**B**) Germination capacity of *afp1-5/2-2/3*-1 mutants seeds obtained from plants cultivated at 21/18 °C after flowering at harvest and during after-ripening at 6 and 12 weeks after harvest (WAH) (Means±SD, n=6 biological replicates, unpaired t-test *P*-value is indicated). (**C**) Germination capacity of single a*fp2-2* and *afp4-2* or double *afp2-2/4-2* mutant seeds at harvest and during after-ripening at 6 and 12 weeks after harvest (WAH) (Means±SD, n=6 biological replicates). (**D**) Germination capacity of freshly harvested Col-0 and *afp1-5/2-2/3-1* seeds in the presence of mock (0.05 % DMSO) or 50 µM of fluridone. (Means±SD, n=6 biological replicates). In (**A**, **C** and **D**), letters on the top of the bars indicate significantly different groups using a one-way ANOVA with a Tukey HSD test (α=0.05)

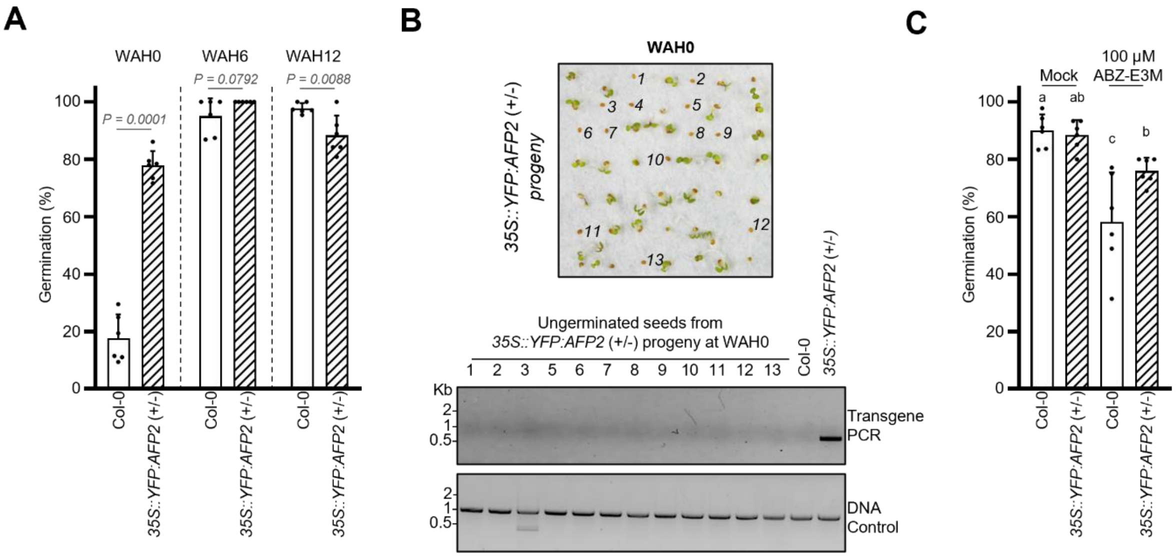


**Fig. S9: AFP2 is a negative regulator of dormancy *sensu stricto*.** (**A**) Germination capacity of seed progenies from *35S::YFP:AFP2 (+/-)* in Col-0 at harvest, and during after-ripening at 6 and 12 weeks after harvest (WAH) (Means±SD, n=6 biological replicates, unpaired t-test *P*-values are indicated). (**B**) Germination assay using progeny of *35S::YFP:AFP2* (+/-) at WAH0 in optimal conditions (top). Subsequently, genomic DNA was isolated from initially non-germinated seeds (labelled from 1 to 13) after their germination was induced through 50 µM GA4+7. (Bottom) Genotyping showing the absence of the *YFP:AFP2* transgene using specific primers for the fusion construct (transgene PCR). Col-0 and *35S::YFP:AFP2* (+/-) seedlings were used as negative and positive amplification controls, respectively. A control for the suitability of DNA preparation for PCR is presented. The seed #4 did not develop into a seedling after stimulation of germination. (**C**) Germination capacity of after-ripened seed progenies from *35S::YFP:AFP2 (+/-)* in Col-0 in presence of mock (0.05 % DMSO) or 100 µM ABZ-E3M (Means±SD, n=6 biological replicates). Letters on the top of the bars indicate significantly different groups using a one-way ANOVA with a Tukey HSD (α=0.05).

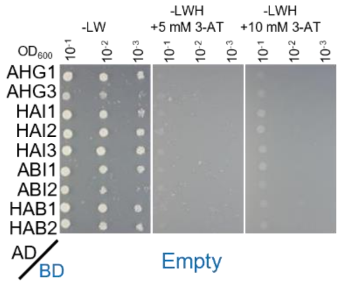


**Fig. S10: Controls for yeast two-hybrid assays.** Yeast was co-transformed with GAL4 AD-PP2CAs and the cognate GAL4 BD-empty vectors and dropped in a serial dilution on control selective growth medium lacking leucine and tryptophan (-LW) or interaction selection medium additionally lacking histidine (-LWH) and containing 3-AT = 3-aminotriazole. No yeast growth on selective -LWH medium was observed for any combination excluding PP2CA DNA-binding activity in this assay. Photos were taken 2 and 4 days after dropping for -LW and -LWH, respectively.

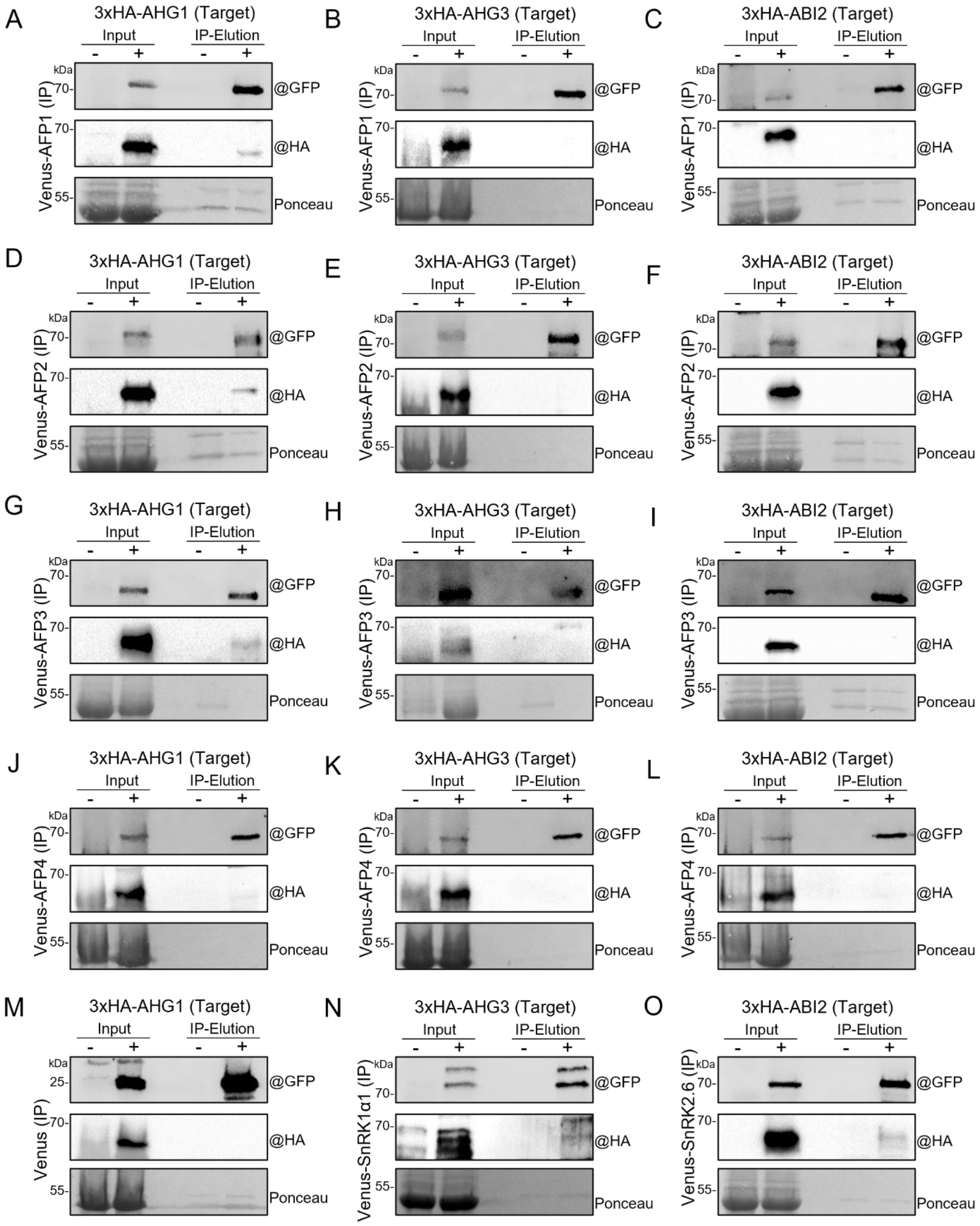
A
3xHA-AHG1 (Target)
Input
IP-Elution
- + - +
kDa
70-
70-
55-
@GFP
@HA
Ponceau
Venus-AFP1 (IP)
B
3xHA-AHG3 (Target)
Input
IP-Elution
- + - +
kDa
70-
70-
55-
@GFP
@HA
Ponceau
Venus-AFP1 (IP)
C
3xHA-ABI2 (Target)
Input
IP-Elution
- + - +
kDa
70-
70-
55-
@GFP
@HA
Ponceau
Venus-AFP1 (IP)
D
3xHA-AHG1 (Target)
Input
IP-Elution
- + - +
kDa
70-
70-
55-
@GFP
@HA
Ponceau
Venus-AFP2 (IP)
E
3xHA-AHG3 (Target)
Input
IP-Elution
- + - +
kDa
70-
70-
55-
@GFP
@HA
Ponceau
Venus-AFP2 (IP)
F
3xHA-ABI2 (Target)
Input
IP-Elution
- + - +
kDa
70-
70-
55-
@GFP
@HA
Ponceau
Venus-AFP2 (IP)
G
3xHA-AHG1 (Target)
Input
IP-Elution
- + - +
kDa
70-
70-
55-
@GFP
@HA
Ponceau
Venus-AFP3 (IP)
H
3xHA-AHG3 (Target)
Input
IP-Elution
- + - +
kDa
70-
70-
55-
@GFP
@HA
Ponceau
Venus-AFP3 (IP)
I
3xHA-ABI2 (Target)
Input
IP-Elution
- + - +
kDa
70-
70-
55-
@GFP
@HA
Ponceau
Venus-AFP3 (IP)
J
3xHA-AHG1 (Target)
Input
IP-Elution
- + - +
kDa
70-
70-
55-
@GFP
@HA
Ponceau
Venus-AFP4 (IP)
K
3xHA-AHG3 (Target)
Input
IP-Elution
- + - +
kDa
70-
70-
55-
@GFP
@HA
Ponceau
Venus-AFP4 (IP)
L
3xHA-ABI2 (Target)
Input
IP-Elution
- + - +
kDa
70-
70-
55-
@GFP
@HA
Ponceau
Venus-AFP4 (IP)
M
3xHA-AHG1 (Target)
Input
IP-Elution
- + - +
kDa
25-
70-
55-
@GFP
@HA
Ponceau
Venus (IP)
N
3xHA-AHG3 (Target)
Input
IP-Elution
- + - +
kDa
70-
70-
55-
@GFP
@HA
Ponceau
Venus-SnRK1α1 (IP)
O
3xHA-ABI2 (Target)
Input
IP-Elution
- + - +
kDa
70-
70-
55-
@GFP
@HA
Ponceau
Venus-SnRK2.6 (IP)

**Fig. S11: AFPs interact specifically with AHG1 *in planta*.** Co-immunoprecipitation (Co-IP) assays in *N. benthamiana* leaves of 3xHA:AHG1, 3xHA:AHG3 and 3xHA:ABI2 (as representative PP2Cs controlled by DOG1, ABA or both, respectively) with Venus:AFP1 (**a-c**), Venus:AFP2 (**D-F**), Venus:AFP3 (**G-I**), and Venus:AFP4 (**J-L**). (**M**) Negative control IP ruling out an interaction of 3xHA:AHG1 with the Venus-tag. (**N&O**) Positive control IPs for 3xHA:AHG3 and 3xHA:ABI2 with known interactors (SnRK1α1 and SnRK2.6, respectively) (*87, 88)*. Non-infiltrated (-) leaves were used aside infiltrated (+) leaves as controls for antibody detection specificities.

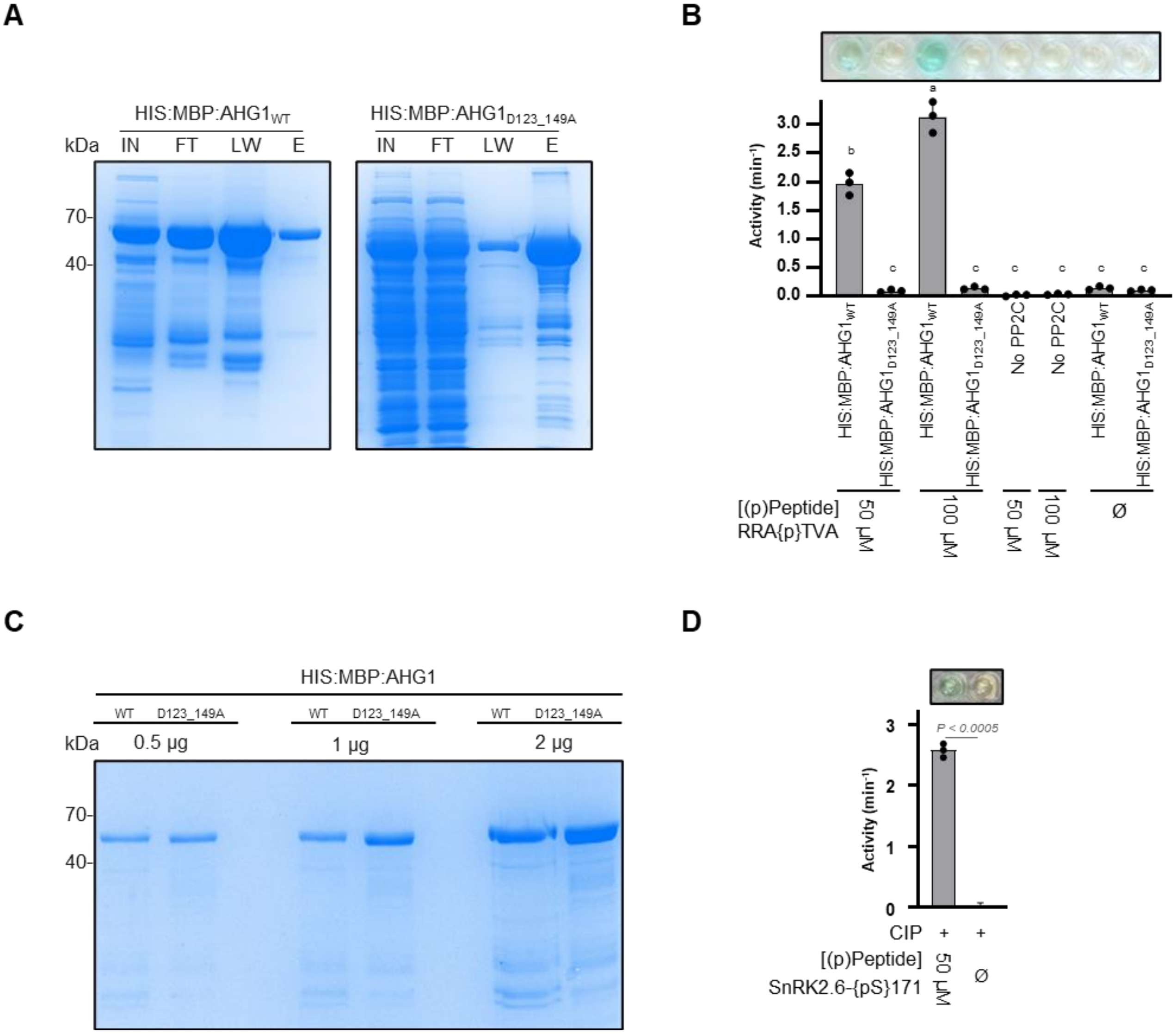
A
HIS:MBP:AHG1WT
HIS:MBP:AHG1D123_149A
kDa
IN
FT
LW
E
70
40
B
Activity (min-1)
3.0
2.5
2.0
1.5
1.0
0.5
0.0
a
b
c
HIS:MBP:AHG1WT
HIS:MBP:AHG1D123_149A
No PP2C
[(p)Peptide]
RRA{p}TVA
50 µM
100 µM
Ø
C
HIS:MBP:AHG1
WT
D123_149A
0.5 µg
1 µg
2 µg
D
3
2
1
0
P < 0.0005
CIP
+
[(p)Peptide]
SnRK2.6-{pS}171

**Fig. S12: Characterization of recombinant HIS:MBP:AHG1 proteins.** (**A**) Coomassie-stained SDS gels showing the purification process for the recombinant HIS:MBP:AHG1 wild type (WT) and mutant D123_149A proteins used for *in vitro* dephosphorylation assays. IN = INPUT, FT = FLOW-THROUGH, LW = LAST WASH FRACTION, E = ELUTION. (**B**) *In vitro* phosphatase assay using 4.4 µM of either recombinant HIS:MBP:AHG1$_{WT}$ or HIS:MBP-AHG1$_{D123_149A}$ and the RRA{p}TVA standard peptide as substrate. Phosphatase activity was measured after 5 minutes of incubation at room temperature. Bar charts show the mean activity and standard deviation of three independent assays for each reaction. Letters on the top of the bars indicate significantly different groups using a one-way ANOVA with a Tukey HSD test ($\alpha$=0.05). (**C**) Coomassie-stained SDS gel directly comparing the loading of HIS:MBP:AHG1$_{WT}$ with HIS:MBP:AHG1$_{D123_149A}$. Different amounts of the purified proteins were loaded. (**D**) *In vitro* phosphatase assays using a commercial calf alkaline intestinal phosphatase (CIP) and the synthetic phosphorylated peptide of SnRK2.6 {pS}171 (KSSVLM{pS}QPKSTVG) corresponding to the phosphorylation site identified in the proteomic analysis. Bar chart showing the mean activity and standard deviation of three independent assays for each reaction, paired t-test *P*-value is indicated. In **B** and **D**, reaction lacking the substrate, or the phosphatase served as negative control and the top panel shows a representative picture of the outcome of the colorimetric assay.

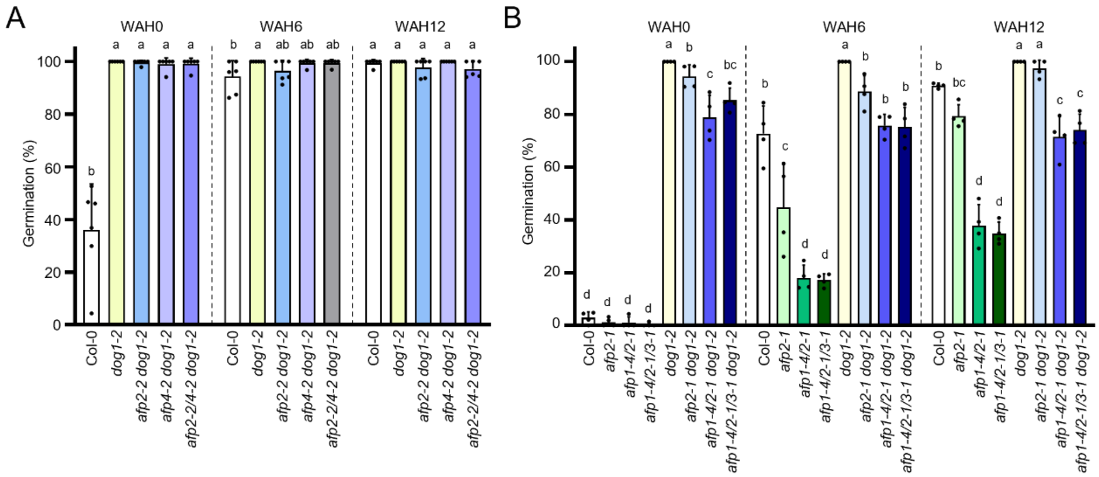


**Fig. S13: Genetic confirmation of the requirements of AFP1 and AFP2 for DOG1 to control dormancy using independent *afp* mutant alleles.** (**A**) Germination capacity of double *afp2-2 dog1-2, afp4-2 dog1-2,* and triple *afp2-2/4-2 dog1-2* at harvest and during after ripening at 6 and 12 weeks after harvest (WAH). (**B**) Germination capacity of single *afp2-1* double *afp1-4/2-1* and triple *afp1-4/2-1/3-1* mutants in a WT or *dog1-2* genetic background at harvest and during after-ripening at 6 and 12 weeks after harvest (WAH). Means±SD, n=6 biological replicates are shown, and letters on the top of the bars indicate significantly different groups using a one-way ANOVA and a Tukey HSD test (α=0.05).

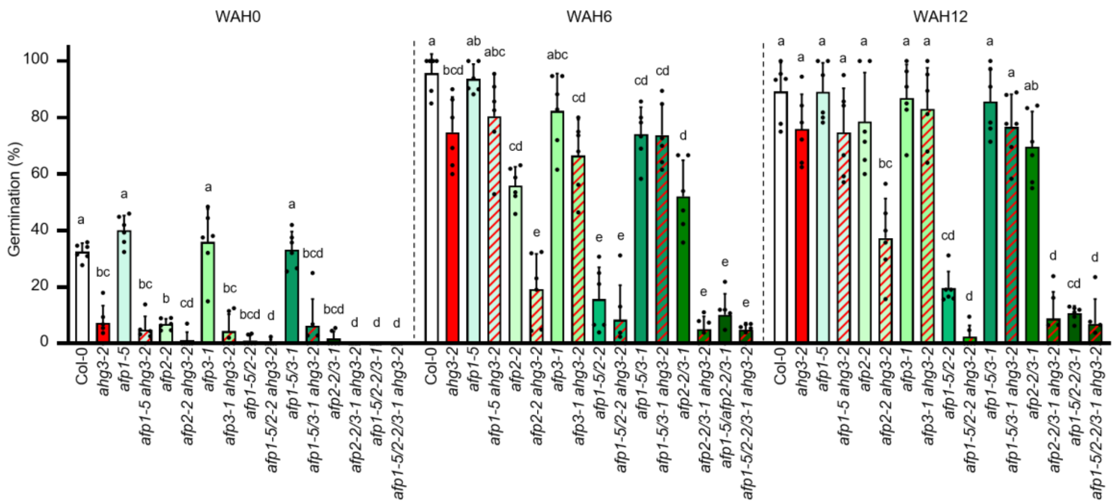


**Fig. S14: Dormancy phenotype of *afp* single and multiple mutants is enhanced by *ahg3-2* mutation.** Germination capacity of *ahg3-2* and *afps* single or combinatorial mutants at harvest and during after-ripening at 6 and 12 weeks after harvest (WAH). Means±SD, n=6 biological replicates are shown, and letters on the top of the bars indicate significantly different groups using a one-way ANOVA and a Tukey HSD test (α=0.05).

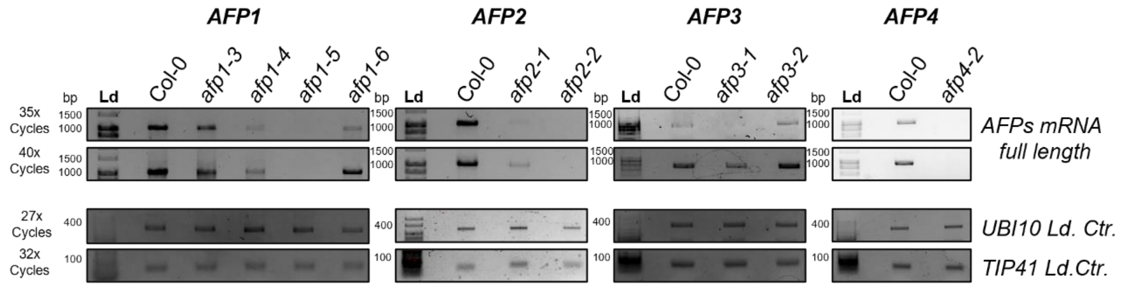


**Fig. S15: Transcript analysis of *afp* mutant alleles.** Semi-Quantitative PCR analysis of *AFP1-4* full length mRNA in dry seeds of the corresponding single *afp* mutants. *TIP41* and *UBI10* amplicons served as loading control.

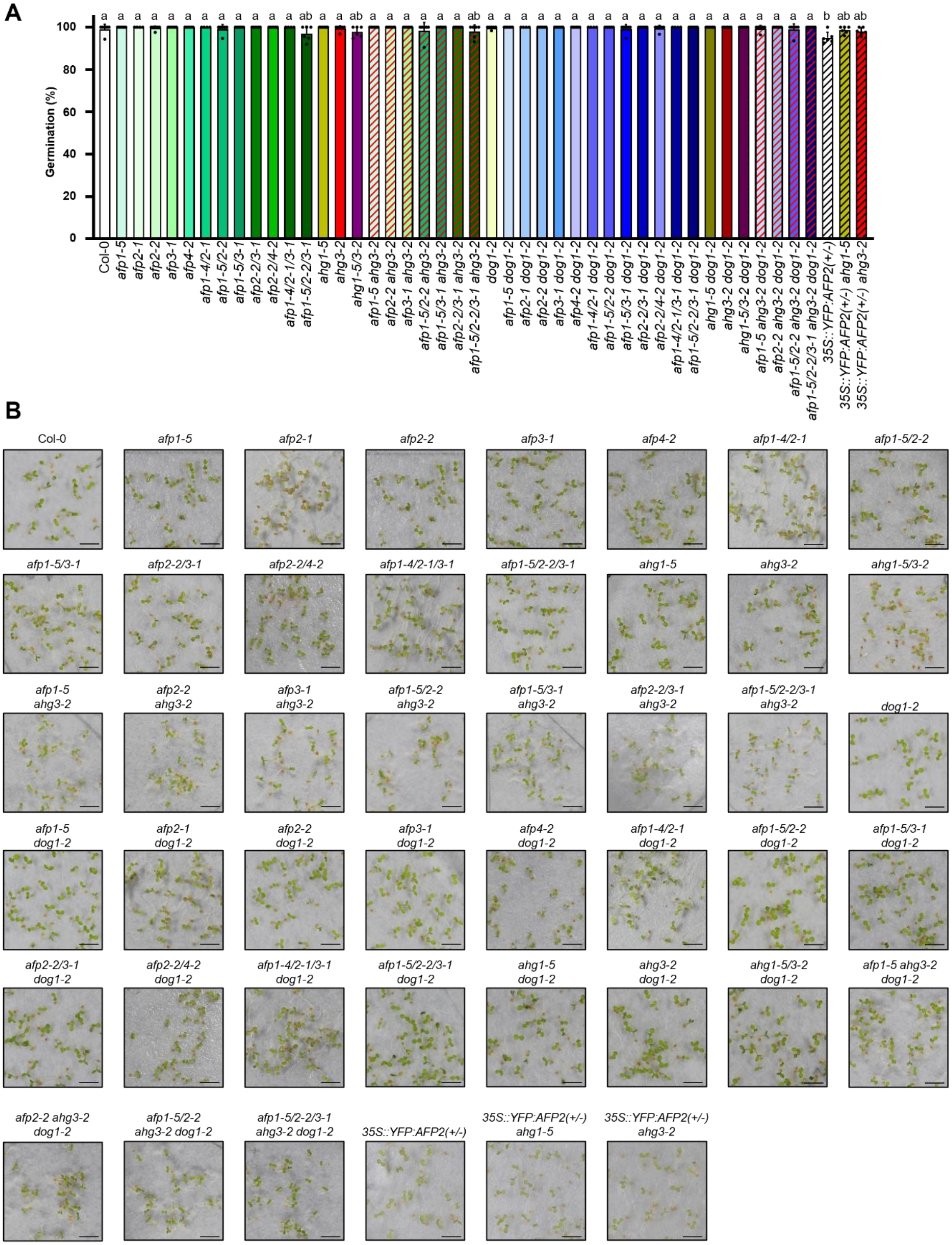
A
Germination (%)
B
Col-0
afp1-5
afp2-1
afp2-2
afp3-1
afp4-2
afp1-4/2-1
afp1-5/2-2
afp1-5/3-1
afp2-2/3-1
afp2-2/4-2
afp1-4/2-1/3-1
afp1-5/2-2/3-1
ahg1-5
ahg3-2
ahg1-5/3-2
afp1-5 ahg3-2
afp2-2 ahg3-2
afp3-1 ahg3-2
afp1-5/2-2 ahg3-2
afp1-5/3-1 ahg3-2
afp2-2/3-1 ahg3-2
afp1-5/2-2/3-1 ahg3-2
dog1-2
afp1-5 dog1-2
afp2-1 dog1-2
afp2-2 dog1-2
afp3-1 dog1-2
afp4-2 dog1-2
afp1-4/2-1 dog1-2
afp1-5/2-2 dog1-2
afp1-5/3-1 dog1-2
afp2-2/3-1 dog1-2
afp2-2/4-2 dog1-2
afp1-4/2-1/3-1 dog1-2
afp1-5/2-2/3-1 dog1-2
ahg1-5 dog1-2
ahg3-2 dog1-2
ahg1-5/3-2 dog1-2
afp1-5 ahg3-2 dog1-2
afp2-2 ahg3-2 dog1-2
afp1-5/2-2 ahg3-2 dog1-2
afp1-5/2-2/3-1 ahg3-2 dog1-2
35S::YFP:AFP2(+/-)
35S::YFP:AFP2(+/-) ahg1-5
35S::YFP:AFP2(+/-) ahg3-2

**Fig. S16: Viability test for seed batches grown at 16/14°C (day/night) temperature regime.** (**A**) Germination capacity of all genotypes used in this study after dormancy alleviation treatment. Seeds were incubated on 50 µM $GA_{4+7}$ and stratified at 4°C for 48 hours before incubation in optimal conditions for Arabidopsis seed germination. Means±SD, n=6 biological replicates are shown, and letters on the top of the bars indicate significantly different groups using a one-way ANOVA and a Tukey HSD test (α=0.05). (**B**) Representative pictures of the assay. Photos were taken 7 days after incubation in optimal conditions for Arabidopsis seed germination and are presented as visual support of the data shown in (**A**). Scale bars represent 2 mm.

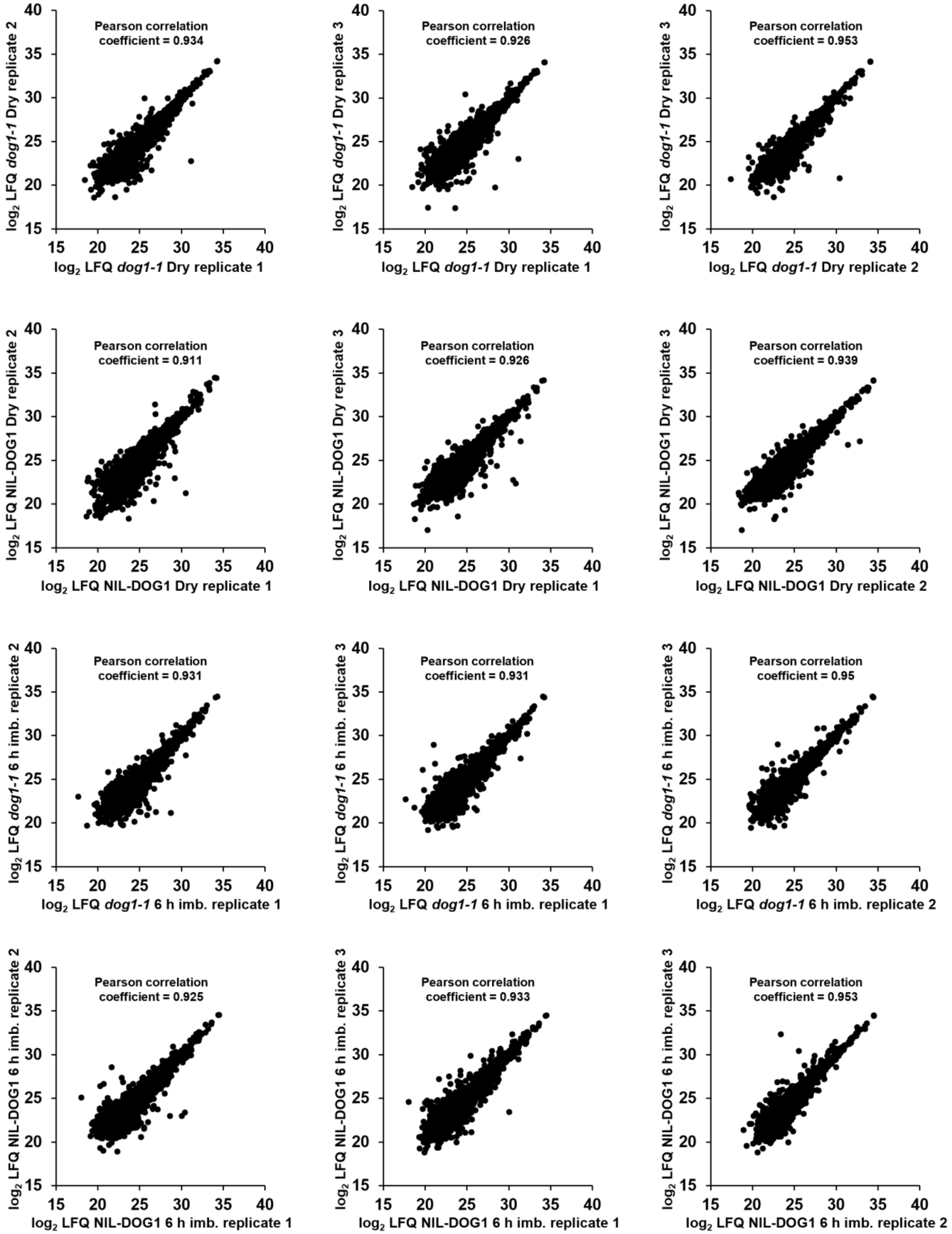
Pearson correlation coefficient = 0.934
log2 LFQ dog1-1 Dry replicate 2
log2 LFQ dog1-1 Dry replicate 1
Pearson correlation coefficient = 0.926
log2 LFQ dog1-1 Dry replicate 3
log2 LFQ dog1-1 Dry replicate 1
Pearson correlation coefficient = 0.953
log2 LFQ dog1-1 Dry replicate 3
log2 LFQ dog1-1 Dry replicate 2
Pearson correlation coefficient = 0.911
log2 LFQ NIL-DOG1 Dry replicate 2
log2 LFQ NIL-DOG1 Dry replicate 1
Pearson correlation coefficient = 0.926
log2 LFQ NIL-DOG1 Dry replicate 3
log2 LFQ NIL-DOG1 Dry replicate 1
Pearson correlation coefficient = 0.939
log2 LFQ NIL-DOG1 Dry replicate 3
log2 LFQ NIL-DOG1 Dry replicate 2
Pearson correlation coefficient = 0.931
log2 LFQ dog1-1 6 h imb. replicate 2
log2 LFQ dog1-1 6 h imb. replicate 1
Pearson correlation coefficient = 0.931
log2 LFQ dog1-1 6 h imb. replicate 3
log2 LFQ dog1-1 6 h imb. replicate 1
Pearson correlation coefficient = 0.95
log2 LFQ dog1-1 6 h imb. replicate 3
log2 LFQ dog1-1 6 h imb. replicate 2
Pearson correlation coefficient = 0.925
log2 LFQ NIL-DOG1 6 h imb. replicate 2
log2 LFQ NIL-DOG1 6 h imb. replicate 1
Pearson correlation coefficient = 0.933
log2 LFQ NIL-DOG1 6 h imb. replicate 3
log2 LFQ NIL-DOG1 6 h imb. replicate 1
Pearson correlation coefficient = 0.953
log2 LFQ NIL-DOG1 6 h imb. replicate 3
log2 LFQ NIL-DOG1 6 h imb. replicate 2

**Fig. S17: Correlation analysis of label free quantification (LFQ) intensities from biological replicates of the total proteomes.** Plotted are $log_2$ transformed LFQ intensities from two replicates, and Pearson correlation coefficients are indicated.

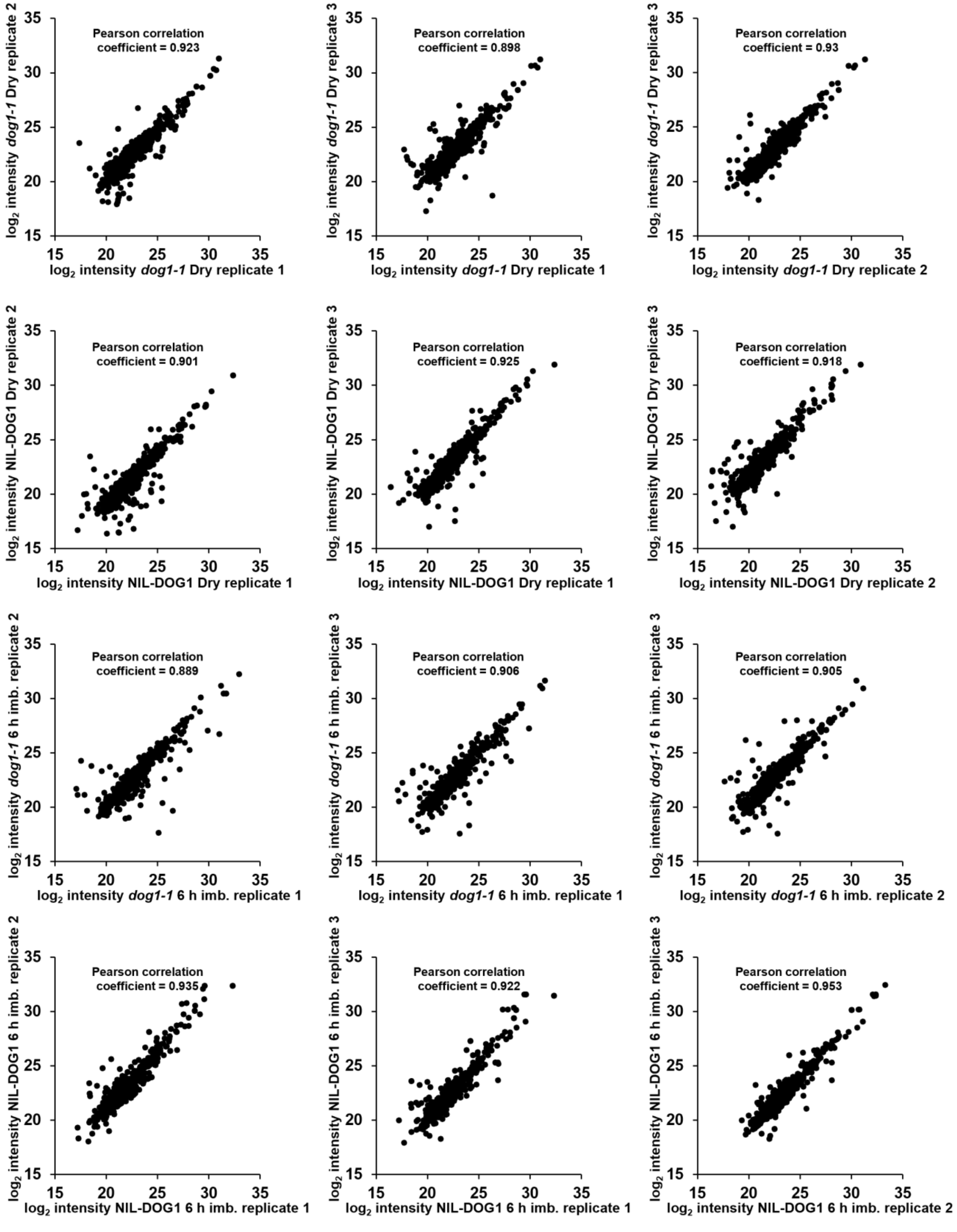

Pearson correlation coefficient = 0.923
log2 intensity dog1-1 Dry replicate 1
log2 intensity dog1-1 Dry replicate 2
Pearson correlation coefficient = 0.898
log2 intensity dog1-1 Dry replicate 1
log2 intensity dog1-1 Dry replicate 3
Pearson correlation coefficient = 0.93
log2 intensity dog1-1 Dry replicate 2
log2 intensity dog1-1 Dry replicate 3
Pearson correlation coefficient = 0.901
log2 intensity NIL-DOG1 Dry replicate 1
log2 intensity NIL-DOG1 Dry replicate 2
Pearson correlation coefficient = 0.925
log2 intensity NIL-DOG1 Dry replicate 1
log2 intensity NIL-DOG1 Dry replicate 3
Pearson correlation coefficient = 0.918
log2 intensity NIL-DOG1 Dry replicate 2
log2 intensity NIL-DOG1 Dry replicate 3
Pearson correlation coefficient = 0.889
log2 intensity dog1-1 6 h imb. replicate 1
log2 intensity dog1-1 6 h imb. replicate 2
Pearson correlation coefficient = 0.906
log2 intensity dog1-1 6 h imb. replicate 1
log2 intensity dog1-1 6 h imb. replicate 3
Pearson correlation coefficient = 0.905
log2 intensity dog1-1 6 h imb. replicate 2
log2 intensity dog1-1 6 h imb. replicate 3
Pearson correlation coefficient = 0.935
log2 intensity NIL-DOG1 6 h imb. replicate 1
log2 intensity NIL-DOG1 6 h imb. replicate 2
Pearson correlation coefficient = 0.922
log2 intensity NIL-DOG1 6 h imb. replicate 1
log2 intensity NIL-DOG1 6 h imb. replicate 3
Pearson correlation coefficient = 0.953
log2 intensity NIL-DOG1 6 h imb. replicate 2
log2 intensity NIL-DOG1 6 h imb. replicate 3

**Fig. S18: Correlation analysis of phosphopeptide intensities from biological replicates.** Plotted are $\log_2$ transformed phospho-peptide intensities from two replicates, and the Pearson correlation coefficients are indicated.

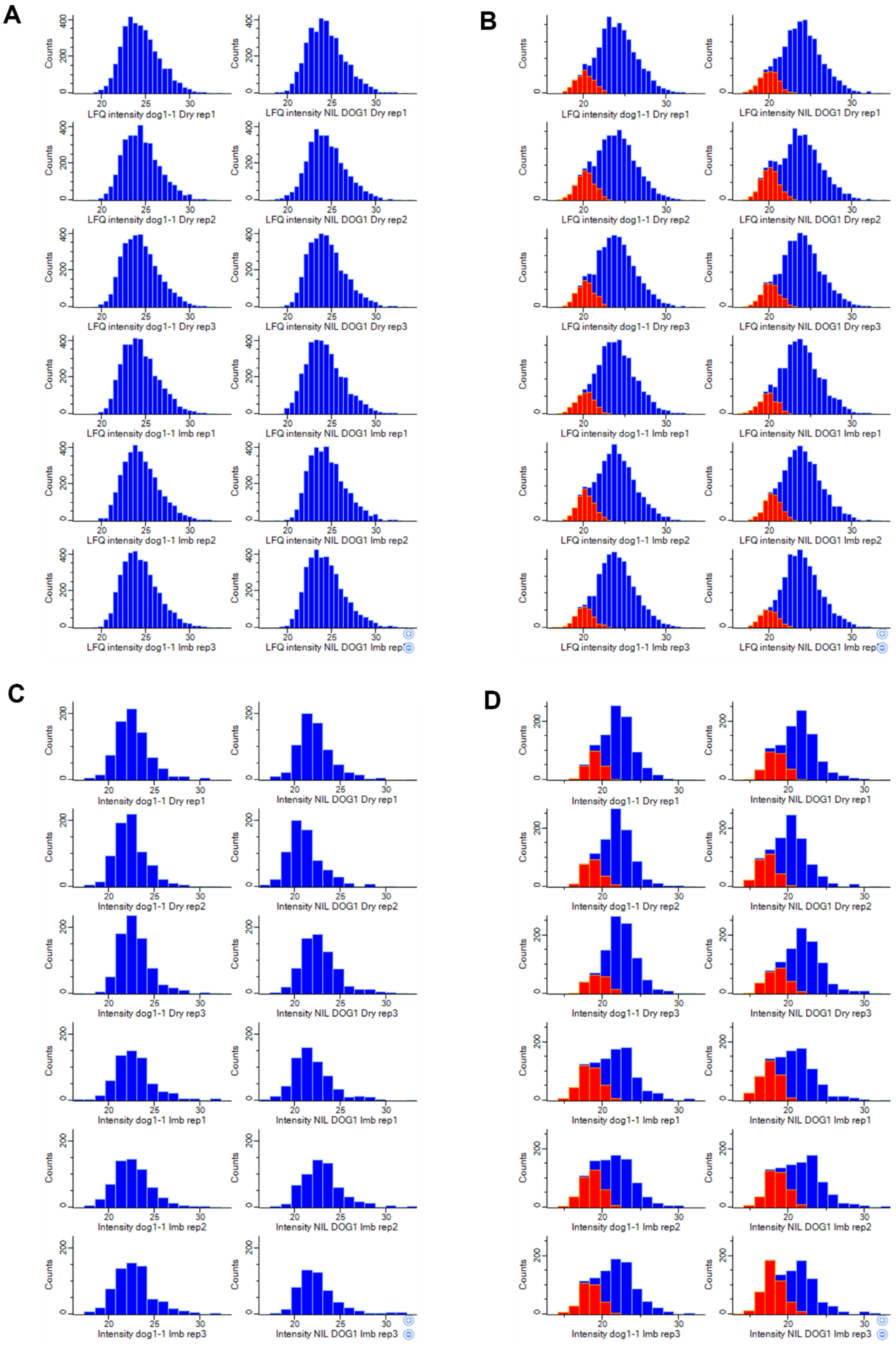
A
Counts
LFQ intensity dog1-1 Dry rep1
LFQ intensity NIL DOG1 Dry rep1
LFQ intensity dog1-1 Dry rep2
LFQ intensity NIL DOG1 Dry rep2
LFQ intensity dog1-1 Dry rep3
LFQ intensity NIL DOG1 Dry rep3
LFQ intensity dog1-1 Imb rep1
LFQ intensity NIL DOG1 Imb rep1
LFQ intensity dog1-1 Imb rep2
LFQ intensity NIL DOG1 Imb rep2
LFQ intensity dog1-1 Imb rep3
B
C
Intensity dog1-1 Dry rep1
Intensity NIL DOG1 Dry rep1
Intensity dog1-1 Dry rep2
Intensity NIL DOG1 Dry rep2
Intensity dog1-1 Dry rep3
Intensity NIL DOG1 Dry rep3
Intensity dog1-1 Imb rep1
Intensity NIL DOG1 Imb rep1
Intensity dog1-1 Imb rep2
Intensity NIL DOG1 Imb rep2
Intensity dog1-1 Imb rep3
Intensity NIL DOG1 Imb rep3
D

**Fig. S19: Distribution of $\log_2$ transformed LFQ (A-B) or phospho-peptide (C-D) intensities across samples.** Histograms show $\log_2$ transformed intensity values before (**A**, **C**) and after (**B, D**) imputation. Values were imputed in Perseus using the imputation from normal distribution setting for each sample with a downshift of 2 or 1.95, and a width of 0.5 or 0.6 for protein groups (**B**) or phosphopeptides (**D**), respectively. Distribution of imputed $\log_2$ transformed LFQ or phospho-peptide intensities are presented as red bars within the histograms shown in panels (**B**) and (**D**).

**Table S1.** General metrics from the MS analysis of the proteome and phosphoproteome of dry and 6 h imbibed NIL-DOG1 and *dog1-1* seeds.

| | **Proteome** | | **Phosphoproteome** | |
|---|---|---|---|---|
| | **Dry seeds** | **6 h imbibed seeds** | **Dry seeds** | **6 h imbibed seeds** |
| Total | 5004 Proteins | | 1616 phospho-sites | |
| Quantified in at least 2 replicates of one genotype | 4324 | | 984 | |
| | 4255 | 4285 | 938 | 859 |
| Phosphoproteins | *#NA* | | 618 | 577 |
| *Exact P-site localization* | *#NA* | | *795* | *734* |
| Regulated in *dog1-1* | 460 | 561 | 186 | 74 |
| *Regulated in dog1-1 exact P-site* | *#NA* | | 144 | 65 |
| Up regulated in *dog1-1* | 265 | 284 | 163 | 59 |
| *Up regulated in dog1-1 (exact P-site)* | *#NA* | | 124 | 51 |
| Down regulated in *dog1-1* | 195 | 277 | 23 | 14 |
| *Down regulated in dog1-1 (exact P-site)* | *#NA* | | 20 | 14 |

**Table S2. Phosphorylated peptides derived of ABA response proteins regulated in seeds of *dog1-1* compared to NIL-DOG1.** Intensity $log_2$ fold change of the indicated p-peptides is listed with an indication of the modified residue and its localization probability. If the residue could not clearly be determined, two positions are listed. For p-peptides that may derive from multiple precursors, SnRK2.6 and ABF1 are indicated as reference sequences to indicate the position of the modified amino acids.

| Protein | AGI code | Residue | Probability (%) | $log_2$ FC | *P*-value | Localization (SUBAcon) | Score | Delta score | Peptide |
|---|---|---|---|---|---|---|---|---|---|
| **OBAP1a** | AT1G05510 | 235 | 100 | 2.71 | 0.015 | cytosol | 102.52 | 87.35 | EVDIKPVE{p}SVPR |
| **UP6** | AT1G16730 | 50 | 96 | -2.78 | 0.026 | peroxisome | 47.52 | 31.92 | ISPESDENVGQGNLY{p}SPDHAIK |
| **STZ** | AT1G27730 | 8; 10 | 50; 50 | 3.27 | 0.016 | nucleus | 57.34 | 43.19 | ALEAL{p}T{p}SPR |
| **ABFs** | AT1G49720 | 81 | 100 | 2.03 | 0.017 | nucleus | 86.88 | 53.45 | QG{p}SLTLPR |
| **AFP1** | AT1G69260 | 115 | 100 | -3.15 | 0.006 | nucleus | 85.61 | 54.00 | TT{p}SLPAESEEEWRK |
| **EM6** | AT2G40170 | 29 | 100 | -1.90 | 0.036 | nucleus | 77.91 | 63.94 | {p}SFEAQQHLAEGR |
| **EEL** | AT2G41070 | 21 | 100 | 1.60 | 0.048 | nucleus | 61.65 | 24.45 | QN{p}SLYSLK |
| **CAP160** | AT4G25580 | 287 | 100 | 1.90 | 0.026 | nucleus | 58.43 | 39.99 | GAGEQP{p}SVAAFGR |
| **PLP4** | AT4G37050 | 9 | 98 | 4.26 | 0.001 | cytosol | 88.18 | 54.22 | GSIS{p}SSEISR |
| **AT5G45690** | AT5G45690 | 4; 5 | 50; 50 | 3.55 | 0.048 | cytosol | 81.49 | 60.79 | A{p}S{p}SDERPGAYPAR |
| **SnRK2IIIs** | AT4G33950 | 171 | 100 | 1.84 | 0.044 | nucleus, cytosol | 75.81 | 62.01 | SSVLH{p}SQPK |
| **ABCC5** | AT1G04120 | 863 | 98 | 2.36 | 0.040 | vacuole | 95.40 | 56.99 | EVQEGG{p}SASDLK |
| **AFP2** | AT1G13740 | 112 | 95 | -2.98 | 0.018 | nucleus | 70.62 | 51.37 | TT{p}SLPAEMEEEWRK |
| **HY5** | AT5G11260 | 36 | 100 | 2.35 | 0.034 | nucleus | 56.35 | 46.37 | EGIE{p}SEEIR |

### Table S3. Primers used for genotyping.

| Name | Sequence (5' – 3') | Purpose |
|---|---|---|
| **afp1-3 LP** | ACCCGTTAGCCAATTGGTAAC | Genotyping of *afp1-3* mutants |
| **afp1-3 RP** | AAATTGACGAGACACGTTTCG | Genotyping of *afp1-3* mutants |
| **afp1-4 LP** | AACCATAAGCCCATTCCTCTG | Genotyping of *afp1-4* mutants |
| **afp1-4 RP** | TCGTTCTCATCAAACCCGTAC | Genotyping of *afp1-4* mutants |
| **afp1-5 LP** | TGTACAGCAGCAAACAAGTGC | Genotyping of *afp1-5* mutants |
| **afp1-5 LP** | AGAGTCGGAGGAAGAGTGGAG | Genotyping of *afp1-5* mutants |
| **afp1-6 LP** | CTCATGTCACAACTGGTGTGG | Genotyping of *afp1-6* mutants |
| **afp1-6 RP** | ACAAAGCCAACAAATCGTTTG | Genotyping of *afp1-6* mutants |
| **afp2-1 LP** | CTCCATCAAACGATTAGTCGC | Genotyping of *afp2-1* mutants |
| **afp2-1 RP** | CCTTTGTTGTCTGCTTGCTTC | Genotyping of *afp2-1* mutants |
| **afp2-2 LP** | ACGACACGTTTCTTGAAGCAG | Genotyping of *afp2-2* mutants |
| **afp2-2 RP** | GATTTTCGGGCTTCTTTCATC | Genotyping of *afp2-2* mutants |
| **afp3-1 LP** | TGTCAGCGATTTTATTTTATTTTG | Genotyping of *afp3-1* mutants |
| **afp3-1 RP** | TCTGTTTCTTCGACATGAGCC | Genotyping of *afp3-1* mutants |
| **afp3-2 LP** | TACCGCCTTAAAAATGAATCG | Genotyping of *afp3-2* mutants |
| **afp3-2 RP** | AGGGGTTTGTTGTATCGGAAC | Genotyping of *afp3-2* mutants |
| **afp4-2 LP** | TGTTGGTAGTCTTGGCTTTGC | Genotyping of *afp4-2* mutants |
| **afp4-2 RP** | GGGTCGATCTTCTTCGATACC | Genotyping of *afp4-2* mutants |
| **ahg1-5 LP** | ACCGACACGTGTTCTGTCTTC | Genotyping of *ahg1-5* mutant |
| **ahg1-5 RP** | CTAAAACTCGACCACCAGCTG | Genotyping of *ahg1-5* mutant |
| **ahg3-2 LP** | TTTGGTTGATTTTAGGTTGCG | Genotyping of *ahg3-2* mutant |
| **ahg3-2 RP** | TTCCCCAGCCTGAATTAAGAG | Genotyping of *ahg3-2* mutant |
| **dog1-2_Mse_F** | TTCTTTAGGCTCGTTTATGCTTTGTGTGGTT | Genotyping of *dog1-2* mutant |
| **dog1-2_Mse_R** | CTGACTACCGAACCAAAAAATTGAATTTAGTC | Genotyping of *dog1-2* mutant |
| **SALK-LB1.3** | ATTTTGCCGATTTCGGAAC | Genotyping of SALK T-DNA lines |
| **AFP2_mid_RW** | TTGCAGCTGTTTGATGATCC | Genotyping of YFP:AFP2 overexpressing lines |
| **YFP_D197FW** | ACAACCACTACCTGAGCTACC | Genotyping of YFP:AFP2 overexpressing lines |
| **SnRK2.6 LP** | GTGAGTGGTCCAATGGATTTG | DNA control PCR |
| **SnRK2.6 RP** | CATATCTTTAGACGAGGGGCC | DNA control PCR |

#### Table S4. Crosses for the obtention of combinatorial high order mutants.

| Mother Plant | Origin of male gametophyte | Genotypes isolated in F2 progenies | Denomination in this study |
|---|---|---|---|
| *afp1-5 (-/-)* | *afp2-2 (-/-)* | *afp1-5 (-/-) afp2-2 (-/-)* | *afp1-5/2-2* |
| *afp1-4 (-/-)* | *afp2-1 (-/-)* | *afp1-4 (-/-) afp2-1 (-/-)* | *afp1-4/2-1* |
| *afp1-5 (-/-) afp2-2 (-/-)* | *afp3-1 (-/-)* | *afp1-5 (-/-) afp3-1 (-/-)*<br>*afp2-2 (-/-) afp3-1 (-/-)*<br>*afp1-5 (-/-) afp2-2 (-/-) afp3-1 (-/-)* | *afp1-5/3-1*<br>*afp2-2/3-1*<br>*afp1-5/2-2/3-1* |
| *afp1-4 (-/-) afp2-1 (-/-)* | *afp3-1 (-/-)* | *afp2-1 (-/-) afp3-1 (-/-)*<br>*afp1-4 (-/-) afp2-1 (-/-) afp3-1 (-/-)* | *afp2-1/3-1*<br>*afp1-4/2-1/3-1* |
| *afp2-2 (-/-)* | *afp4-2 (-/-)* | *afp2-2 (-/-) afp4-2 (-/-)* | *afp2-2/4-2* |
| *dog1-2 (-/-)* | *afp1-5 (-/-)* | *afp1-5 (-/-) dog1-2 (-/-)* | *afp1-5 dog1-2* |
| *dog1-2 (-/-)* | *afp2-2 (-/-)* | *afp2-2 (-/-) dog1-2 (-/-)* | *afp2-2 dog1-2* |
| *dog1-2 (-/-)* | *afp2-1 (-/-)* | *afp2-1 (-/-) dog1-2 (-/-)* | *afp2-1 dog1-2* |
| *dog1-2 (-/-)* | *afp3-1 (-/-)* | *afp3-1 (-/-) dog1-2 (-/-)* | *afp3-1 dog1-2* |
| *dog1-2 (-/-)* | *afp1-5 (-/-) afp2-2 (-/-)* | *afp1-5 (-/-) afp2-2 (-/-) dog1-2 (-/-)* | *afp1-5/2-2 dog1-2* |
| *dog1-2 (-/-)* | *afp1-4 (-/-) afp2-1 (-/-)* | *afp1-4 (-/-) afp2-1 (-/-) dog1-2 (-/-)* | *afp1-4/2-1 dog1-2* |
| *afp1-5 (-/-) afp2-2 (-/-) dog1-2 (-/-)* | *afp3-1 (-/-) dog1-2 (-/-)* | *afp1-5 (-/-) afp3-1 (-/-) dog1-2 (-/-)*<br>*afp2-2 (-/-) afp3-1 (-/-) dog1-2 (-/-)*<br>*afp1-5 (-/-) afp2-2 (-/-) afp3-1 (-/-) dog1-2 (-/-)* | *afp2-2/3-1 dog1-2*<br>*afp2-2/3-1 dog1-2*<br>*afp1-5/2-2/3-1 dog1-2* |
| *afp1-4 (-/-) afp2-1 (-/-) dog1-2 (-/-)* | *afp3-1 (-/-) dog1-2 (-/-)* | *afp1-4 (-/-) afp2-1 (-/-) afp3-1 (-/-) dog1-2(-/-)* | *afp1-4/afp2-1/afp3-1 dog1-2* |
| *afp1-5 (-/-) afp2-2 (-/-) afp3-1 (-/-)* | *ahg3-2 (-/-)* | *afp1-5 (-/-) ahg3-2 (-/-)*<br>*afp2-2 (-/-) ahg3-2 (-/-)*<br>*afp3-1 (-/-) ahg3-2 (-/-)*<br>*afp1-5 (-/-) afp2-2 (-/-) ahg3-2 (-/-)*<br>*afp1-5 (-/-) afp3-1 (-/-) ahg3-2 (-/-)*<br>*afp2-2 (-/-) afp3-1 (-/-) ahg3-2 (-/-)*<br>*afp1-5 (-/-) afp2-2 (-/-) afp3-1 (-/-) ahg3-2 (-/-)* | *afp1-5 ahg3-2*<br>*afp2-2 ahg3-2*<br>*afp3-1 ahg3-2*<br>*afp1-5/3-1 ahg3-2*<br>*afp1-5/afp3-1 ahg3-2*<br>*afp2-2/3-1 ahg3-2*<br>*afp1-5/2-2/3-1 ahg3-2* |
| *afp1-5 (-/-) afp2-2 (-/-) dog1-2 (-/-)* | *ahg3-2 (-/-) dog1-2 (-/-)* | *afp1-5 (-/-) ahg3-2 (-/-) dog1-2 (-/-)*<br>*afp2-2 (-/-) afp3-1 (-/-) ahg3-2 (-/-) dog1-2 (-/-)*<br>*afp1-5 (-/-) afp2-2 (-/-) ahg3-2 (-/-) dog1-2 (-/-)*<br>*afp1-5 (-/-) afp2-2 (-/-) afp3-1 (-/-)ahg3-2 (-/-) dog1-2 (-/-)* | *afp1-5 ahg3-2 dog1-2*<br>*afp3-1 ahg3-2 dog1-2*<br>*afp1-5/2-2 ahg3-2 dog1-2*<br>*afp1-5/2-2/3-1 ahg3-2 dog1-2* |
| *35S::YFP:AFP2(+/-)* | *ahg1-5 (-/-)* | *35S::YFP:AFP2(+/-)ahg1-5 (-/-)* | *35S::YFP:AFP2(+/-) ahg1-5* |
| *35S::YFP:AFP2(+/-)* | *ahg3-2 (-/-)* | *35S::YFP:AFP2(+/- )ahg3-2 (-/-)* | *35S::YFP:AFP2(+/-) ahg3-2* |

### Table S5. Primers used for transcript analysis.

| Name | Sequence (5' – 3') | Purpose |
|---|---|---|
| **AFP1CDS_FW** | ATGGCGGAAGCAAACGAG | Semi Q-PCR analysis of full-length *AFP1* |
| **AFP1CDS_RW** | GGTCTATTATAAGAGATTAGAAGG | Semi Q-PCR analysis of full-length *AFP1* |
| **AFP2CDS_FW** | AAGCAAGCAGACAACAAAGG | Semi Q-PCR analysis of full-length *AFP2* |
| **AFP2CDS_RW** | CGATTTAAAAGGTCGAAGAAGAGG | Semi Q-PCR analysis of full-length *AFP2* |
| **AFP3CDS_FW** | ATGTCGAAGAAACAGAGATTGTCTG | Semi Q-PCR analysis of full-length *AFP3* |
| **AFP3CDS_RW** | TCACAAGAAGGGAGATGGATTTAC | Semi Q-PCR analysis of full-length *AFP3* |
| **AFP4CDS_FW** | ATGGAGATGATAAGAGGCA | Semi Q-PCR analysis of full-length *AFP4* |
| **AFP4CDS_RW** | CTAGATAGAAGAAGAAGGAAGTTTA | Semi Q-PCR analysis of full-length *AFP4* |
| **UBI10sqPCR_FW** | CTCTCTACCGTGATCAAGATGCAGATC | Semi Q-PCR analysis of UBI10 as reference gene |
| **UBI10sqPCR_RW** | GGATCTTGGCCTTAACGTTGTCG | Semi Q-PCR analysis of UBI10 as reference gene |
| **TIP41sqPCR_FW** | TCGTGAAAACTGTTGGAGAGAAGCAA | Semi Q-PCR analysis of TIP41 as reference gene |
| **TIP41sqPCR_RW** | TCAACTGGATACCCTTTCGCA | Semi Q-PCR analysis of TIP41 as reference gene |
| **RCAR1qPCR_FW** | TTCCTCAAGGTAACACAAAGGA | Q-PCR analysis of RCAR1 |
| **RCAR1qPCR_RW** | CACTTCAATGCCCTTGTTCA | Q-PCR analysis of RCAR1 |
| **RCAR2qPCR_FW** | TCTGAAAGATTGGCTGCTCA | Q-PCR analysis of RCAR2 |
| **RCAR2qPCR_RW** | GCATGATCACACGTGCTTCT | Q-PCR analysis of RCAR2 |
| **RCAR3qPCR_FW** | GAACGTCTTGCGGTTCAAG | Q-PCR analysis of RCAR3 |
| **RCAR3qPCR_RW** | TTTTTGCCTTGTCCATCCTC | Q-PCR analysis of RCAR3 |
| **RCAR4qPCR_FW** | GCTGGAGGTTGGTAGCGTAA | Q-PCR analysis of RCAR4 |
| **RCAR4qPCR_RW** | CCTTTGTGTTACCTTCCGGC | Q-PCR analysis of RCAR4 |
| **RCAR5qPCR_FW** | TGCCGGAAGGAAATAGTGAG | Q-PCR analysis of RCAR5 |
| **RCAR5qPCR_RW** | ATTTCCCATCGACCATGACT | Q-PCR analysis of RCAR5 |
| **RCAR6qPCR_FW** | TGTGGAGAGTTATGTGGTGGA | Q-PCR analysis of RCAR6 |
| **RCAR6qPCR_RW** | CAATCCCACGACTTTTACAACTT | Q-PCR analysis of RCAR6 |
| **RCAR7qPCR_FW** | GTCGAGACCATTGAAGCACCA | Q-PCR analysis of RCAR7 |
| **RCAR7qPCR_RW** | GGAAGCCGGAGACTAACGT | Q-PCR analysis of RCAR7 |
| **RCAR8qPCR_FW** | ATCGGTGACGACACTACACG | Q-PCR analysis of RCAR8 |
| **RCAR8qPCR_RW** | ATGAGATTATTGCCGGTTGG | Q-PCR analysis of RCAR8 |
| **RCAR9qPCR_FW** | CACGTCATCAGTTTCAGCGT | Q-PCR analysis of RCAR9 |
| **RCAR9qPCR_RW** | GTGTTCTCGGCGAGTTTAGC | Q-PCR analysis of RCAR9 |
| **RCAR10qPCR_FW** | ACGTCGTTGATGTTCCTCCA | Q-PCR analysis of RCAR10 |
| **RCAR10qPCR_RW** | ACATAGATCACACGACGGCT | Q-PCR analysis of RCAR10 |
| **RCAR11qPCR_FW** | CGTAACTCCGGTGACGGAAG | Q-PCR analysis of RCAR11 |
| **RCAR11qPCR_RW** | AACCTTGCACGTCATTCTCA | Q-PCR analysis of RCAR11 |
| **RCAR12qPCR_FW** | CGGAAGGTAATTCGGAGGA | Q-PCR analysis of RCAR12 |
| **RCAR12qPCR_RW** | CCCCCAAATTTCATCATTACC | Q-PCR analysis of RCAR12 |
| **RCAR13qPCR_FW** | TCAACGAGTTCGTCGTCTTG | Q-PCR analysis of RCAR13 |
| **RCAR13qPCR_RW** | CACTTTCAGGTCGGAGAAGC | Q-PCR analysis of RCAR13 |
| **RCAR14qPCR_FW** | TCTAAGCTTCAGGGTCGTCG | Q-PCR analysis of RCAR14 |
| **RCAR14qPCR_RW** | TTCCTCTGTGTTTCCCTCGG | Q-PCR analysis of RCAR14 |
| **AFP1qPCR_FW** | ACGGCAGTGGAGAAGAAGTT | Q-PCR analysis of AFP1 |
| **AFP1qPCR_RW** | GGTTGGTTAGAGGGTTTTTGG | Q-PCR analysis of AFP1 |
| **AFP2qPCR_FW** | TGTGCATTTGCCATGGTAGT | Q-PCR analysis of AFP2 |
| **AFP2qPCR_RW** | ACAAGACCCTTCCGGTTTCT | Q-PCR analysis of AFP2 |
| **AFP3qPCR_FW** | GCAAAGAAGCGAGCCAAAAC | Q-PCR analysis of AFP3 |
| **AFP3qPCR_RW** | AATTCTGCCGGTGAGAGGAA | Q-PCR analysis of AFP3 |
| **AFP4qPCR_FW** | CGATGATGATGGTGGTGATG | Q-PCR analysis of AFP4 |
| **AFP4qPCR_RW** | GCTTCAGCTAGTTTAGTCGGTCA | Q-PCR analysis of AFP4 |
| **PEX4_qPCR_FW** | TTACGAAGGCGGTGTTTTTC | Q-PCR analysis of PEX4 as reference gene |
| **PEX4_qPCR_RW** | GGCGAGGCGTGTATACATTT | Q-PCR analysis of PEX4 as reference gene |

**Table S6. Primary antibodies used in this study.** All antibodies were diluted in PBS buffer at 0.1 % tween 20.

| Antibody | Target | Host | Dilution | Reference |
|---|---|---|---|---|
| *@SnRK2IIIs* | SnRK2.2 (AT3G50500)<br>SnRK2.3 (AT5G66880)<br>SnRK2.6 (AT4G33950) | Rabbit | 1:3000 | Agrisera AS14 2783 |
| *@AFP1* | AFP1 (AT1G69260) | Rabbit | 1:2000 | (*49*) |
| *@ABI5* | ABI5 (AT2G36270) | Rabbit | 1:2500 | Abcam ab98831 |
| *@H3* | Histones H3 variants | Rabbit | 1:15000 | Abcam ab1791 |
| *@Actin* | Actin variants | Rabbit | 1:10000 | Sigma A-2066 |
| *@GFP* | GFP_AEQVI and variants including Venus | Rabbit | 1:4000 | Abcam ab290 |
| *@HA* | Influenza hemagglutinin-HA-epitope | Rabbit | 1:7500 | Abcam ab9110 |

**Table S7. Secondary antibodies used in this study.** All antibodies were diluted in PBS buffer at 0.1 % tween 20.

| Antibody | Target | Host | Deatection methods | Dilution | Reference |
|---|---|---|---|---|---|
| *@Rabbit ECL* | Rabbit IgG H&L | Goat | ECL-Chemidoc Imager | 1:25000 | Agrisera AS09 602 |
| *@Rabbit IR* | Rabbit IgG H&L | Goat | IR800 channel Odyssey Imager | 1:35000 | LI-COR IRDye® 800CW |

#### **Table S8.** Primers used for cloning and site-directed mutagenesis.

| Name | Sequence (5' – 3') | Purpose |
|---|---|---|
| **AFP1attext_FW** | AAAAAGCAGGCTATATGGCGGAAGCAAACGAG | Cloning of *AFP1* with attB sites |
| **AFP1attext_RW** | AGAAAGCTGGGTATTAGAAGGAGAAGAAGTGTTGACAAC | Cloning of *AFP1* with attB sites |
| **AFP2attext_FW** | AAAAAGCAGGCTATATGGGTGAAGCAAGCAG | Cloning of *AFP2* with attB sites |
| **AFP2attext_RW** | AGAAAGCTGGGTAATTCTTCGATTTAGAAGGTCG | Cloning of *AFP2* with attB sites |
| **AFP3attext_FW** | AAAAAGCAGGCTATATGTCGAAGAAACAGAGATTGTCTG | Cloning of *AFP3* with attB sites |
| **AFP3attext_RW** | AGAAAGCTGGGTATCACGCGACTGCAAGCTTT | Cloning of *AFP3* with attB sites |
| **AFP4attext_FW** | AAAAAGCAGGCTATATGGAGATGATAAGAGGCA | Cloning of *AFP4* with attB sites |
| **AFP4attext_RW** | AGAAAGCTGGGTACTAGATAGAAGAAGAAGGAAGTTTA | Cloning of *AFP4* with attB sites |
| **ABI1attext_FW** | AAAAAGCAGGCTATATGGAGGAAGTATCTCCGGC | Cloning of *ABI1* with attB sites |
| **ABI1attext_RW** | AGAAAGCTGGGTATCAGTTCAAGGGTTTGCTCTTG | Cloning of *ABI1* with attB sites |
| **HAB1attext_FW** | AAAAAGCAGGCTATATGGAGGAGATGACTCCCG | Cloning of *HAB1* with attB sites |
| **HAB1attext_RW** | AGAAAGCTGGGTATCAGGTTCTGGTCTTGAACTTTC | Cloning of *HAB1* with attB sites |
| **HAB2attext_FW** | AAAAAGCAGGCTATATGGAAGAGATTTCACCTGC | Cloning of *HAB2* with attB sites |
| **HAB2attext_RW** | AGAAAGCTGGGTATCAAGATCTGGTCTTGAACTTTC | Cloning of *HAB2* with attB sites |
| **HAI1attext_FW** | AAAAAGCAGGCTATATGGCTGAGATTTGTTACGAG | Cloning of *HAI1* with attB sites |
| **HAI1attext_RW** | AGAAAGCTGGGTACTACGTGTCTCGTCGTAGATCAAC | Cloning of *HAI1* with attB sites |
| **HAI2attext_FW** | AAAAAGCAGGCTATATGGCGGATATTTGTTATGAAG | Cloning of *HAI2* with attB sites |
| **HAI2attext_RW** | AGAAAGCTGGGTATCAAGCAACGTGTCTCTTTCTTC | Cloning of *HAI2* with attB sites |
| **HAI3attext_FW** | AAAAAGCAGGCTATATGGCCGAGATATGTTACGAAG | Cloning of *HAI3* with attB sites |
| **HAI3attext_RW** | AGAAAGCTGGGTATTATCTTCTGAGATCAATCACAACG | Cloning of *HAI3* with attB sites |
| **SnRK2.6attext_FW** | AAAAAGCAGGCTATATGGATCGACCAGCAGTGAG | Cloning of *SnRK2.6* with attB sites |
| **SnRK2.6attext_RW** | AGAAAGCTGGGTATCACATTGCGTACACAATCTCTC | Cloning of *SnRK2.6* with attB sites |
| **SnRK1a1for attext** | AAAAAGCAGGCTATATGTTCAAACGAGTAGATGAGTT | Cloning of *SnRK1a1* with attB sites |
| **SnRK1a1rev attext** | AGAAAGCTGGGTATCAGAGGACTCGGAGCT | Cloning of *SnRK1a1* with attB sites |
| **attBext for** | ACAAGTTTGTACAAAAAAGCAGGCT | Generation of complete attB sites |
| **attBext rev** | ACCACTTTGTACAAGAAAGCTGGGT | Generation of complete attB sites |
| **AHG1 S149A fw** | CTTTGCGGTATACGCTGGCCACGGCGGTTC | Site-directed mutagenesis of AHG1 |
| **AHG1 S149A rev** | GAACCGCCGTGGCCAGCGTATACCGCAAAG | Site-directed mutagenesis of AHG1 |
| **AHG1 S123A fw** | GATCTCGTAAGATGGAGGCTTCGGTTACTGTTAAACC | Site-directed mutagenesis of AHG1 |
| **AHG1 S123A rev** | GGTTTAACAGTAACCGAAGCCTCCATCTTACGAGATC | Site-directed mutagenesis of AHG1 |

#### **Table S9. Peptides targeted for PRM MS measurements.**

| Protein | AGI code | Sequence | Modification | Charge (z) | Mass/charge (m/z) | Polarity |
|---|---|---|---|---|---|---|
| **SnRK2.6** | AT4G33950 | SSVLHSQPK | No | 2 | 491.769448 | positive |
| **SnRK2.6** | AT4G33950 | SSVLHSQPK (+ $HPO_3$) | (STY) Phosphorylation | 2 | 531.752614 | positive |
| **SnRK2.6** | AT4G33950 | STVGTPAYIAPEVLLK | No | 2 | 829.971813 | positive |
| **SnRK2.6** | AT4G33950 | STVGTPAYIAPEVLLK (+ $HPO_3$) | (STY) Phosphorylation | 2 | 869.955 | positive |
| **SnRK2.6** | AT4G33950 | STVGTPAYIAPEVLLKK | No | 3 | 596.348622 | positive |
| **SnRK2.6** | AT4G33950 | STVGTPAYIAPEVLLKK (+ $HPO_3$) | (STY) Phosphorylation | 3 | 623.0041 | positive |
| **SnRK2.6** | AT4G33950 | STVGTPAYIAPEVLLKK (+2x $HPO_3$) | 2x (STY) Phosphorylation | 3 | 649.6595 | positive |
| **SnRK2.6** | AT4G33950 | ICDFGYSKSSVLHSQPK | No | 2 | 976.980375 | positive |
| **SnRK2.6** | AT4G33950 | ICDFGYSKSSVLHSQPK (+ $HPO_3$) | (STY) Phosphorylation | 2 | 1016.9635 | positive |
| **SnRK2.6** | AT4G33950 | ICDFGYSKSSVLHSQPK | No | 3 | 651.656 | positive |
| **SnRK2.6** | AT4G33950 | ICDFGYSKSSVLHSQPK (+ $HPO_3$) | (STY) Phosphorylation | 3 | 678.3115 | positive |
